# Reactive astrocytes promote proteostasis in Huntington’s disease through the JAK2-STAT3 pathway

**DOI:** 10.1101/2021.04.29.441924

**Authors:** Laurene Abjean, Lucile Ben Haim, Miriam Riquelme-Perez, Pauline Gipchtein, Céline Derbois, Marie-Ange Palomares, Fanny Petit, Anne-Sophie Hérard, Marie-Claude Gaillard, Martine Guillermier, Mylène Gaudin-Guérif, Gwenaelle Aurégan, Nisrine Sagar, Cameron Héry, Noëlle Dufour, Noémie Robil, Mehdi Kabani, Ronald Melki, Pierre De la Grange, Alexis P. Bemelmans, Gilles Bonvento, Jean-François Deleuze, Philippe Hantraye, Julien Flament, Eric Bonnet, Solène Brohard, Robert Olaso, Emmanuel Brouillet, Maria-Angeles Carrillo-de Sauvage, Carole Escartin

**Affiliations:** Université Paris-Saclay, Commissariat à l’Energie Atomique et aux Energies Alternatives, Centre National de la Recherche Scientifique, MIRCen, Laboratoire des Maladies Neurodégénératives, 92265, Fontenay-aux-Roses, France.; Université Paris-Saclay, Commissariat à l’Energie Atomique et aux Energies Alternatives, Centre National de Recherche en Génomique Humaine, 91057, Evry, France.; GenoSplice technology, 75013 Paris, France

**Author notes:** Correspondence to: Carole Escartin, Ph.D. UMR9199 - MIRCen CNRS - CEA - Université Paris Saclay 18, route du Panorama 92260 Fontenay-aux-roses. These authors contributed equally to this work (Lucile Ben Haim, Miriam Riquelme-Perez). Abbreviations: AAV = Adeno-Associated Virus; DARPP32 = Dopamine- and cAMP-Regulated neuronal Phosphoprotein; BSA = Bovine Serum Albumin; DNAJB1 = DnaJ heat shock protein family (Hsp40) member B1; FACS = Fluorescence-Activated Cell Sorting; GFAP = Glial Fibrillary Acidic Protein; Glutamate Chemical Exchange Saturation transfer = gluCEST; HBSS = Hank’s Balanced Salt Solution; HSP = Heat Shock Proteins; JAK = Janus Kinase; LV = Lentiviral Vector; mHTT = mutant Huntingtin; RT = Room Temperature; SDS = Sodium Dodecyl Sulfate; SOCS3 = Suppressor Of Cytokine Signalling 3; STAT3 = Signal Transducer and Activator of Transcription 3; Tx = Triton X-100; Ub = Ubiquitin; UPS = Ubiquitin-Proteasome System; WT = Wild type.

**Keywords:** Neuron-astrocyte interactions, JAK2-STAT3 signalling, Aggregated protein clearance, Viral vectors, Neurodegenerative diseases.

## Abstract

Huntington’s disease is a fatal neurodegenerative disease characterized by striatal neurodegeneration, aggregation of mutant Huntingtin and the presence of reactive astrocytes. Astrocytes are important partners for neurons and engage in a specific reactive response in Huntington’s disease that involves morphological, molecular and functional changes. How reactive astrocytes contribute to Huntington’s disease is still an open question, especially because their reactive state is poorly reproduced in experimental mouse models.

Here, we show that the JAK2-STAT3 pathway, a central cascade controlling astrocyte reactive response, is activated in the putamen of Huntington’s disease patients. Selective activation of this cascade in astrocytes through viral gene transfer reduces the number and size of mutant Huntingtin aggregates in neurons and improves neuronal defects in two complementary mouse models of Huntington’s disease. It also reduces striatal atrophy and increases glutamate levels, two central clinical outcomes measured by non-invasive magnetic resonance imaging. Moreover, astrocyte-specific transcriptomic analysis shows that activation of the JAK2-STAT3 pathway in astrocytes coordinates a transcriptional program that increases their intrinsic proteolytic capacity, through the lysosomal and ubiquitin-proteasome degradation systems. This pathway also enhances their production and exosomal release of the co-chaperone DNAJB1, which contributes to mutant Huntingtin clearance in neurons.

Together, our results show that the JAK2-STAT3 pathway controls a beneficial proteostasis response in reactive astrocytes in Huntington’s disease, which involves bi-directional signalling with neurons to reduce mutant Huntingtin aggregation, eventually improving disease outcomes.

## Introduction

Huntington’s disease is a genetic neurodegenerative disease that causes involuntary movements, psychiatric symptoms and cognitive deficits, with no curative treatment yet available^1^. Huntington’s disease is caused by an expansion of CAG triplet repeats in the *Huntingtin* (*HTT*) gene, leading to an abnormal polyglutamine tract in the N-terminal part of the protein HTT^2^. Both loss and gain of function of HTT contribute to the dysfunction and death of projection neurons in the caudate/putamen (striatum in mice) and cerebral cortex. A key neuropathological hallmark of Huntington’s disease is the presence of mutant HTT (mHTT) aggregates, primarily in neurons^3^, but also in glial cells^4, 5^. In the brain of Huntington’s disease patients, astrocytes progressively become reactive, which was initially characterized by their hypertrophic morphology and overexpression of Glial Fibrillary Acidic Protein (GFAP)^6–8^. Reactive astrocytes also display significant changes in gene expression in the putamen^9^ and cortex^5^ of Huntington’s disease patients. Astrocytes are essential partners of neurons, as they perform many key functions including metabolic and trophic support, antioxidant defence and regulation of synaptic transmission and plasticity^10^. How are these functions changed in Huntington’s disease? Most studies report defective astrocyte functions in Huntington’s disease models^11^, including reduced glutamate uptake^12^, altered K^+^ buffering^13^, impaired regulation of blood flow^14^, as well as reduced synthesis and release of antioxidants^15^, trophic factors^16^, gliotransmitters^17^ and exosomes^18^. However, in most cases, Huntington’s disease mouse models poorly replicate the reactive state of astrocytes observed in Huntington’s disease human brains, based on GFAP overexpression, hypertrophy^11, 13^ and transcriptional profile, as recently captured with genome-wide transcriptomic analysis^9, 19, 20^. Therefore, the impact of reactive astrocytes on disease progression is still unclear.

The Janus Kinase (JAK)-Signal Transducer and Activator of Transcription 3 (STAT3) pathway is a central cascade controlling astrocyte reactive response^21^. STAT3 activation is found in reactive astrocytes of genetic models of Huntington’s disease in mice and non-human primates ^22^. However, it is still unknown whether this pathway is also activated in reactive astrocytes observed in Huntington’s disease patients and which astrocyte functions are regulated by this pathway.

We previously reported that inhibition of the JAK2-STAT3 pathway in reactive astrocytes in an acute model of Huntington’s disease reduces their reactive features and increases the number of mHTT aggregates^22^. mHTT aggregates are mainly composed of N-terminal fragments of mHTT, which trap several important proteins such as transcription factors or chaperones^23, 24^ and generate deleterious steric hindrance. However, aggregates may also contribute to remove toxic soluble mHTT from the cytosol^25^. Soluble mHTT can be degraded by the Ubiquitin-Proteasome System (UPS), while aggregates can only be cleared by autophagy coupled to lysosomal degradation^26, 27^. Astrocytes are reported to have high proteolytic capacity, including for mHTT, which could explain why fewer aggregates are observed in astrocytes than in neurons^28, 29^. Yet, it is unknown whether the UPS and autophagy/lysosome systems are specifically altered in Huntington’s disease astrocytes^30, 31^, and how these systems can be stimulated in astrocytes to promote mHTT clearance. Another important proteostasis mechanism preventing mHTT aggregation is operated by chaperones, which promote HTT proper folding, prevent abnormal interactions with cellular proteins and guide mHTT to degradation systems^32, 33^. In particular, Heat Shock Proteins (HSP) prevent mHTT aggregation in different cell types^32, 34^.

Here, we studied how the JAK2-STAT3 pathway controls astrocyte reactive response in Huntington’s disease, focusing on their ability to promote cellular proteostasis. We observed STAT3 activation in reactive astrocytes in the brain of Huntington’s disease patients. In two complementary mouse models of Huntington’s disease, we found that activation of the JAK2- STAT3 pathway in reactive astrocytes reduces both the number and size of mHTT aggregates in neurons, without increasing soluble mHTT levels, and improves several disease hallmarks. Genome-wide transcriptomics and functional analysis showed that JAK2-STAT3 pathway activation induces a specific proteostasis signature in astrocytes associated with higher proteolytic activity. It also induces astrocyte expression of the co-chaperone DNAJB1, which is loaded in exosomes and reduces mHTT aggregation in neurons. Our results show that the JAK2-STAT3 pathway controls a bi-directional communication between reactive astrocytes and neurons in Huntington’s disease, which eventually reduces mHTT aggregation and improves neuronal alterations.

## Materials and methods

### Mice

Heterozygous knock-in mice (Hdh140 mice) expressing a chimeric mouse/human exon 1 containing 140 CAG repeats inserted into the murine *Htt* gene on a C57BL/6J background were originally obtained from the Jackson laboratory (stock # 027409)^35^. Male and female Hdh140 mice and their littermate controls were injected with different viral vectors (see below) at 7-10 months and euthanized 4 months later. Wild type (WT), adult male C57BL/6J mice were injected at 2.5 months of age, with different viral vectors and euthanized 1.5 months later.

All experimental protocols were reviewed and approved by the local ethics committee (CETEA N°44) and the French Ministry of Education and Research (Approval APAFIS#4554- 2016031709173407). They were performed in an authorized animal facility (authorization #D92-032-02), in strict accordance with recommendations of the European Union (2010- 63/EEC), and in compliance with the 3R guidelines. Animal care was supervised by a dedicated veterinarian and animal technicians. Mice were housed in groups of 5, under standard environmental conditions (12-hour light-dark cycle, temperature: 22 ± 1°C and humidity: 50%) in filtered cages, with *ad libitum* access to food and water.

### Viral vectors

We either used lentiviral vectors (LV) or adeno-associated vectors (AAV) to drive transgene expression in neurons or in astrocytes, as described in each figure. Self-inactivated LV were produced by transient co-transfection of 293T cells with four plasmids encoding viral structural proteins, the envelope protein and the transgenic cDNA under the control of a phosphoglycerate kinase 1 promoter and followed by the woodchuck hepatitis virus post-transcriptional regulatory element, to enhance transgene expression^36^. To target neurons, LV were pseudotyped with the G-protein of the vesicular stomatitis virus^37^. To target astrocytes, LV were pseudotyped with the rabies-related Mokola envelope and lentiviral recombinant genome contained four repeats of the miR124 target to repress transgene expression in neurons^36^. LV are referred to as “LVA- or LVN-name of the transgene” depending the cell type targeted: A for astrocytes and N for neurons. Lentiviral particle titers were determined by ELISA quantification of the nucleocapsid p24 protein.

AAV (AAV2, serotype 9) contain the gfaABC1D promoter, a synthetic promoter derived from the GFAP promoter^38^, and transduce astrocytes. AAV were produced according to validated procedures^39^. Viral genome concentration was determined by qPCR on DNase resistant particles.

Viral vectors encoding murine *Socs3* or *Jak2^T875N^*, a constitutively active form of JAK2 (JAK2ca), were used to respectively, inhibit or activate the JAK2-STAT3 pathway in mouse astrocytes^22, 39, 40^. They were co-injected with a viral vector encoding the fluorescent proteins *Gfp* or *Td-Tomato* to visualize transduced cells (same total viral load). For some experiments, the same viral vectors were injected in both striata. For other experiments, each striatum of the same mouse received different viruses and data were analysed with paired statistical tests.

LVs encoding the first 171 N-terminal amino acids of human *Huntingtin* (*HTT*) cDNA with 82 polyglutamine repeats that target either striatal neurons [LVN-mHTT^41^] or astrocytes [LVA- mHTT^8^] were used as LV-based models of Huntington’s disease.

Last, we generated LVA expressing full-length human *DNAJB1* (LVA-DNAJB1) or the dominant negative mutant corresponding to the J-domain of human *DNAJB1*^42^ [LVA-DNAJB1- DN]. The initial cDNA was generated by GeneArt Gene Synthesis services (Invitrogen, Carlsbad, CA) based on published sequences, and inserted into a pENTR^®^ transfer plasmid. Gateway^®^ recombination (Thermofisher Scientific) was used to clone these cDNA into appropriate self-inactivated LV expression plasmid containing the phosphoglycerate kinase 1 promoter, the woodchuck hepatitis virus post-transcriptional regulatory element and four miR124 target sequences.

### Immunostaining quantification

Levels of GFAP immunoreactivity and GFAP^+^ volumes were quantified on 10x-tiled images of serial sections along the antero-posterior axis of the striatum, acquired with an epifluorescence microscope (Leica DM6000, Nussloch, Germany). Virally transduced GFP^+^ area was manually segmented on each section and the corresponding mean intensity signal for GFAP in this area was extracted with Image J. Background intensity signal was measured on unstained regions of the same section and subtracted to the GFAP total signal. The volume was calculated from the area measured on each section by the Cavalieri method^40^.

To quantify lesion size, images of DARPP32 immunostained serial striatal sections were acquired at the 5x objective with an epifluorescence microscope (Leica DM6000). DARPP32- depleted area in the striatum was manually segmented with Image J on each serial section, and the total volume calculated with the Cavalieri method.

To quantify astrocyte soma area and STAT3 immunoreactivity, stacked confocal images of GFP and STAT3 immunostained sections were acquired with a 40× objective (three brain sections per mouse, three fields per section, 10 to 16 z-steps of 1 μm, maximum intensity stack). GFP^+^ cell bodies were manually segmented and their individual area and mean grey value for STAT3 were measured with Image J.

The total number and surface of EM48^+^ and Ubiquitin^+^ aggregates were quantified on serial striatal sections along the antero-posterior axis, scanned with an Axio scanZ.1 (Zeiss, Oberkochen, Germany) at the 40× objective in bright field microscopy mode, with multi-plan focusing. Aggregates were automatically detected by the Morphostrider software (ExploraNova), with intensity, size and shape thresholds, after manual segmentation of the striatum on each section. The total number of aggregates within each striatum was then calculated. To quantify the distribution of EM48^+^ aggregates in DARPP32^+^ neurons and GFP^+^ astrocytes, stacked confocal images (16 z-steps of 1 µm, maximum intensity stack) were acquired on a Leica TCS SP8 confocal fluorescent microscope with a 40x objective (three brain sections per mouse, three fields per section). Aggregates were automatically detected with ImageJ software, with intensity, size and shape thresholds. Laser intensity, detection settings and analysis parameters were identical between each mouse of the same cohort. The number of aggregates in each cell type and the total number were manually quantified using ImageJ cell counter plugin.

### Quantification of cathepsin and proteasome activities in astrocytes

Hdh140 mice previously injected with astrocyte-specific AAV-GFP or AAV-Td-Tomato in the striatum to label astrocytes, were perfused with cold PBS for 4 min. Their striata were rapidly collected in Hank’s Balanced Salt Solution (HBSS; Sigma). Cells were mechanically and enzymatically dissociated with fire-polished Pasteur pipettes and the neural tissue dissociation kit with papain (Miltenyi Biotec, Bergisch Gladbach, Germany), following manufacturer’s instructions. After filtration through a 50 μm-filter, myelin removal beads II and MS columns (Miltenyi Biotec) were used to deplete myelin from cell suspensions. Cells were then resuspended in 0.5% PNB buffer (Perkin Elmer, FP1020) and incubated for 30 min with 1 µM fluorescent cathepsin probe (iABP, Vergent Bioscience, Minneapolis, MN, #40200) or with 200 nM proteasome probe (UbiQ, Bio BV, Amsterdam, the Netherlands, #UbiQ-018) at RT. Cells were centrifuged at 300 g for 5 min at 4°C and resuspended in 400 µl HBSS. They were sorted on a BD Influx cell sorter. GFP expressed by infected astrocytes was detected at 530/40 nm (488 nm excitation) and the cathepsin probe was detected at 670/30 nm (646 nm excitation). Td-Tomato expressed by infected astrocytes was detected at 579/34 nm (561 nm excitation) and the proteasome probe at 530/40 nm (488 nm excitation). Control samples of unlabelled or mono-fluorescent brain cells were used to define detector gains and sorting gates, which were kept constant for all samples. No compensation was required to accurately quantify the two fluorescent signals within the same cell. Cells were gated on a side scatter/ forward scatter plot, then singlets were selected and finally the percentage of GFP^+^/Cathepsin^+^ or Td- Tomato^+^/Proteasome^+^ astrocytes was quantified in each mouse, after setting the gates on GFP^+^ or Td-Tomato^+^ astrocytes incubated without a probe (i.e. “Fluorescence Minus One” controls, **Supplementary Fig. 1B**).

### Statistics

Values for individual samples are shown on graphs. Arithmetic means are represented by a horizontal line and paired samples from two groups are connected by a line. Sample size was chosen based on prior experience to yield adequate power to detect specific effects. Mice of the appropriate genotype were randomly allocated to experimental groups and processed alternatively. Statistical analysis was performed with Statistica software (StatSoft, Tulsa, OK) and graphs were prepared with GraphPad Prism 7 (La Jolla, CA). Paired or unpaired two-tailed Student *t-*test were used to compare two groups, or two-way ANOVA to compare four groups. Normality of residues and homoscedasticity were assessed. If any condition of application was not fulfilled, we used non-parametric tests: two groups were compared by the Mann-Whitney or Wilcoxon matched-pair tests. Percentages were first changed to proportions and transformed by the *arcsine* function, before being analysed by paired or unpaired *t*-test.

Investigators were partially blinded to the group when performing experiments and analysis, as the group can be deduced by the presence of aggregates or GFP levels for example. The significance level was set at *p* < 0.05. In each figure legend, N refers to the number of mice.

### Data availability

Microarray datasets are deposited on GEO under reference GSE107486. RNAseq datasets are deposited on GEO under reference GSE171141.

Detailed protocols for stereotactic injections, immunohistology, protein extraction, exosome isolation, immunoblot, RNA extraction and RT-qPCR, microarray, RNAseq analysis and identification of transcription factors as well as magnetic resonance imaging are provided as **Supplementary material.**

## Results

### STAT3 activation in reactive astrocytes in Huntington’s disease patients

STAT3 is involved in the control of reactive astrocytes in multiple CNS diseases and corresponding animal models^21^, but it has never been studied in the brain of Huntington’s disease patients. We performed STAT3 immunostainings in the putamen of Huntington’s disease patients (Vonsattel grade III^6^) and their age- and sex-matched controls. There was a stronger STAT3 immunoreactivity in Huntington’s disease patients (**Fig. 1A, B**), especially in regions showing neurodegeneration, as seen by their lower density in NeuN^+^ neurons and higher density in hypertrophic GFAP^+^ astrocytes (**Fig. 1C**). Many STAT3^+^ cells had a typical astrocyte morphology and displayed nuclear accumulation of this transcription factor (**Fig. 1B, C**), an indication of pathway activation^21^.

**Figure 1.**
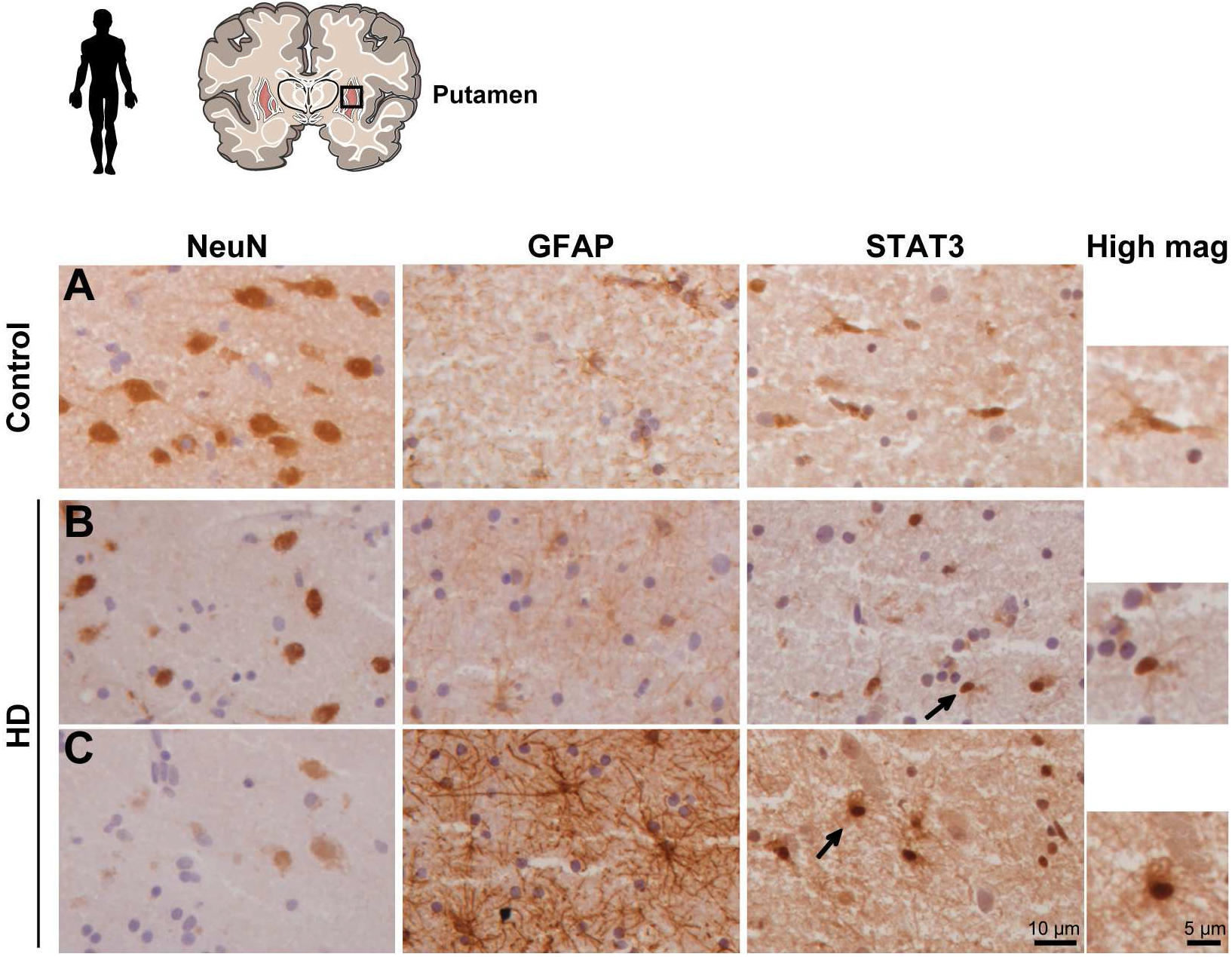
STAT3 nuclear accumulation in the putamen of Huntington’s disease patients. STAT3 immunoreactivity is higher in the putamen of Huntington’s disease patients (**B, C**) than in control subjects (**A**), especially in regions displaying many hypertrophic GFAP^+^ astrocytes, and major neurodegeneration, as seen with the loss NeuN staining (**C**). STAT3 is often found accumulated in the nucleus of cells with a typical astrocyte morphology (arrows and high magnification in **B** and **C**). Representative images from 4 subjects/group.

We also analysed available transcriptomic data of nuclei isolated from the cingulate cortex of grade III/IV Huntington’s disease patients^5^ to identify potential active transcription factors in Huntington’s disease astrocytes. Several bioinformatics tools (based on literature mining or DNA motif recognition in the promoter of differentially expressed genes, see Methods) identified STAT3 as a potential regulatory transcription factor in Huntington’s disease astrocytes, with significant associated *p* values (**Supplementary table 1**).

Together, these results support a role for STAT3 in driving astrocyte reactive changes in Huntington’s disease.

### Astrocytic JAK2-STAT3 pathway reduces neuronal mHTT aggregation in two mouse models of Huntington’s disease

To determine the molecular and functional regulation operated by STAT3 in Huntington’s disease reactive astrocytes, we took advantage of our viral vectors that transduce striatal astrocytes with high efficiency and selectivity (**Supplementary Fig. 1A)**, to either activate or inhibit the JAK2-STAT3 pathway^39, 40^.

We first studied knock-in Hdh140 mice that express a humanized *HTT* gene with 140 CAG repeats under its own endogenous promoter^35^. These mice develop progressive Huntington’s disease symptoms with small intra-neuronal mHTT aggregates, early transcriptional defects in neurons, but very mild morphological and molecular reactive changes in astrocytes^20, 43^. In this model, we thus stimulated the JAK2-STAT3 pathway in striatal astrocytes by virus-mediated expression of a constitutively active form of JAK2 (JAK2ca).

A lentiviral vector targeting astrocytes and encoding JAK2ca (LVA-JAK2ca) was injected in 7-9 month-old heterozygous Hdh140 mice, with a LVA encoding GFP (LVA-GFP) to visualize infected astrocytes (Hdh140-JAK2ca mice, **Fig. 2A**). Controls were Hdh140 mice injected with LVA-GFP at the same total viral titer (Hdh140-GFP mice), and brains from both groups were analysed 4 months later (**Fig. 2A**). Immunostaining on mouse brain sections showed that JAK2ca activated STAT3 (**Fig. 2B**) and induced the two cardinal features of reactive astrocytes: overexpression of the intermediate filaments GFAP (**Fig. 2B, C**) and soma hypertrophy (**Fig. 2B, D**). JAK2ca also increased mRNA levels of *Vimentin* and *Serpina3n,* two markers of reactive astrocytes (**Fig. 2E, F**). We controlled that JAK2ca did not impact total mRNA levels of murine *Htt* and *mHTT,* which were expressed at similar levels in Hdh140- JAK2ca and Hdh140-GFP groups (Student *t-*test, *n* = 3-4/group, *df* = 5, *t* = 1.367*, p* = 0.230 and *t* = 1.276*, p* = 0.258 respectively, data not shown).

**Figure 2.**
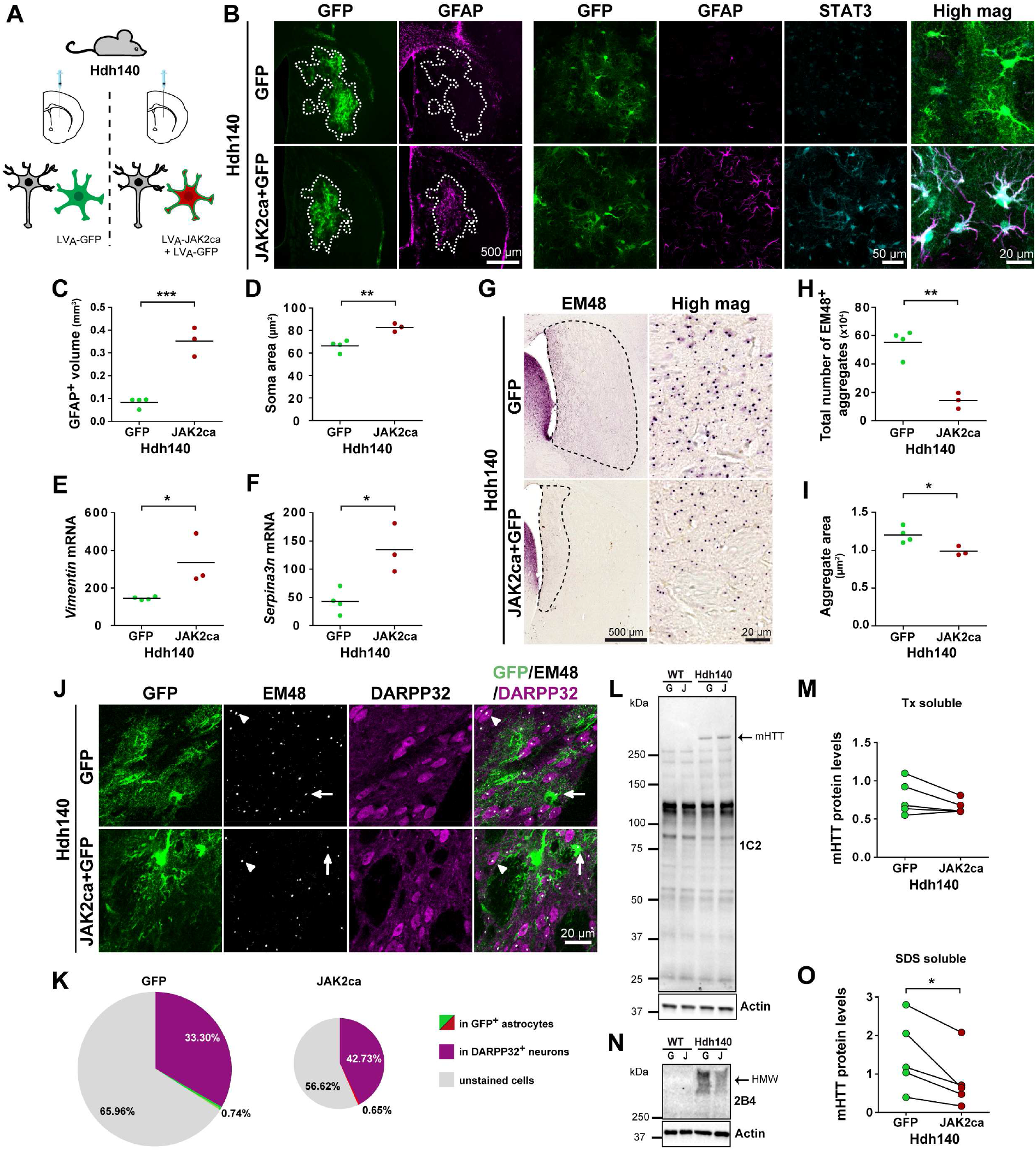
JAK2ca induces astrocyte reactivity and reduces neuronal mHTT aggregation in Hdh140 mice A. Hdh140 mice (7-9 month-old) were injected in the striatum with LVA-GFP or LVA-JAK2ca + LVA-GFP, at the same total virus load, and their brains analysed 4 months later. **B.** Low magnification images (left) show the GFP^+^ transduced area (outlined, green) and GFAP staining (magenta) in the striatum of Hdh140-GFP and Hdh140-JAK2ca mice. High magnification images (right) of astrocytes stained for GFP (green), GFAP (magenta), and STAT3 (cyan). JAK2ca triggers STAT3 activation, as seen by its nuclear accumulation, increases GFAP and vimentin levels and induces morphological changes in astrocytes. Note that the basal expression of GFAP is nearly undetectable in Hdh140 mice, suggesting very mild reactive changes in this model. **C, D.** GFAP^+^ volume (**C**) and soma area of GFP^+^ astrocytes (**D**) are significantly increased by JAK2ca. Student *t*-test. *n* = 3-5/group, *df* = 5, **C**: *t* = 8.081, *p* = 0.0005; **D**: *t* = 4.703, *p* = 0.0053. **E, F.** JAK2ca increases *Vimentin* (**E**) and *Serpina3n* (**F**) mRNA levels. Student *t*-test. *n* = 3-5/group, *df* = 5, **E**: *t* = 2.917, *p* = 0.0331; **F**: *t* = 3.748, *p* = 0.0133. **G.** Bright field images of EM48^+^ aggregates in the striatum of Hdh140-GFP and Hdh140-JAK2ca mice. The striatal region displaying EM48^+^ aggregates is outlined on low magnification images. **H, I.** Total number (**H**) and size (**I**) of EM48^+^ aggregates are significantly decreased by JAK2ca in the striatum of Hdh140 mice. Student *t*-test. *n* = 3-5/group, *df* = 5, **H**: *t* = 6.606, *p* = 0.0012; **I**: *t* = 3.142, *p* = 0.0256. **J.** Confocal images of striatal sections stained for GFP (green), EM48 (white) and DARPP32 (magenta). EM48^+^ aggregates are mostly found in neurons labelled with DARPP32 (arrowhead) and very rarely in GFP^+^ astrocytes (green, arrow). **K**. JAK2ca decreases the total number of EM48^+^ aggregates, but the distribution of EM48^+^ aggregates between GFP^+^ astrocytes and DARPP32^+^ neurons is not changed. **L, M.** Immunoblotting on the Tx-soluble fraction of mHTT with the 1C2 antibody. Similar levels of mHTT are detected in Hdh140-GFP and Hdh140-JAK2ca mice, while mHTT is undetectable in WT mice. There is no mHTT cleavage fragments detected by this antibody in both Hdh140-GFP and Hdh140-JAK2ca mice. Paired *t*-test, *n* = 5, *df* = 4, *t* = 1.845 *p* = 0.139. **N, O.** Immunoblotting on the SDS-soluble fraction of high molecular weight species (HMW) of mHTT with the 2B4 antibody. JAK2ca decreases the levels of insoluble HMW mHTT species in Hdh140 mice. Band intensity was normalized to actin. Paired *t*-test. *n* = 5/group, *df* = 4, *t* = 3.367, *p* = 0.0281.

JAK2ca expression in Hdh140 astrocytes significantly reduced the total number of mHTT aggregates in the striatum (**Fig. 2G, H**). The size of EM48^+^ aggregates was also significantly decreased in Hdh140-JAK2ca mice (**Fig. 2I**). Most EM48^+^ aggregates were in small neuronal processes, while more than 30% of them were found in the nucleus or cell body of striatal neurons expressing dopamine- and cAMP-regulated neuronal phosphoprotein (DARPP32), and less than 1% of all mHTT aggregates were found in GFP^+^ astrocytes (**Fig. 2J, K**). Despite the marked reduction in the number of mHTT aggregates, this relative distribution was not significantly impacted by JAK2ca (Student *t-*test on *arcsine-*transformed data, *n* = 3-4/group, *df* = 5, *t* = 1.983*, p* = 0.104 for DARPP32^+^ neurons, *t* = 0.517*, p* = 0.627 for GFP^+^ astrocytes, **Fig. 2K**).

To determine whether soluble mHTT was also reduced by JAK2ca, we analysed Triton X- 100 (Tx)-soluble protein extracts prepared from the striatum of WT and Hdh140 mice injected with LVA-GFP or LVA-JAK2ca. Immunoblotting with the 1C2 antibody that recognizes preferentially the elongated polyglutamine stretch of mHTT^44^ showed similar Tx-soluble mHTT levels in both groups (**Fig. 2L, M**). We then immunoblotted proteins from the Tx-insoluble, SDS-soluble fraction with the 2B4 antibody, which preferentially binds to the N-terminal part of mHTT^45^. High molecular weight forms of mHTT corresponding to aggregated fragments of mHTT were detected only in samples from Hdh140 mice. The levels of high molecular weight aggregated mHTT were lower in the Hdh140-JAK2ca group, in accordance with histological data of reduced mHTT aggregation with JAK2ca (**Fig. 2N, O**).

We then took advantage of another lentiviral mouse model of Huntington’s disease that better replicates the strong neurodegeneration and subsequent astrocyte reactivity observed in patients^8, 22^. This model involves lentivirus-mediated expression of the 171 first amino acids of human HTT with 82 CAG repeats specifically in striatal neurons^8, 41^ (**Fig. 3A**). In this model, we performed the inverse manipulation of the JAK2-STAT3 pathway in astrocytes, by blocking it with its inhibitor Suppressor Of Cytokine Signalling 3 (SOCS3). SOCS3 efficiently blocked STAT3 activation and reactive changes in astrocytes (**Fig. 3B-F**)^22^. Moreover, SOCS3 expression in striatal astrocytes increased the aggregation of mHTT in neurons, as seen with EM48 immunostaining (**Fig. 3G, H**). Less than 2% of mHTT aggregates were found in GFP^+^ astrocytes (**Fig. 3H**), and this distribution was not changed by SOCS3 (Student *t-*test on *arcsine*- transformed data, *n* = 6/group, *df* = 5, *t* = 0.917*, p* = 0.401). mHTT aggregates are ubiquitinated and the total number of Ubiquitin (Ub)^+^ aggregates was also increased by SOCS3 (**Fig. 3I, J**). In addition, aggregates were larger in the SOCS3 group than in the GFP group (**Fig. 3K**), revealing both quantitative and qualitative changes in neuronal mHTT aggregates following JAK2-STAT3 inhibition in astrocytes.

**Figure 3.**
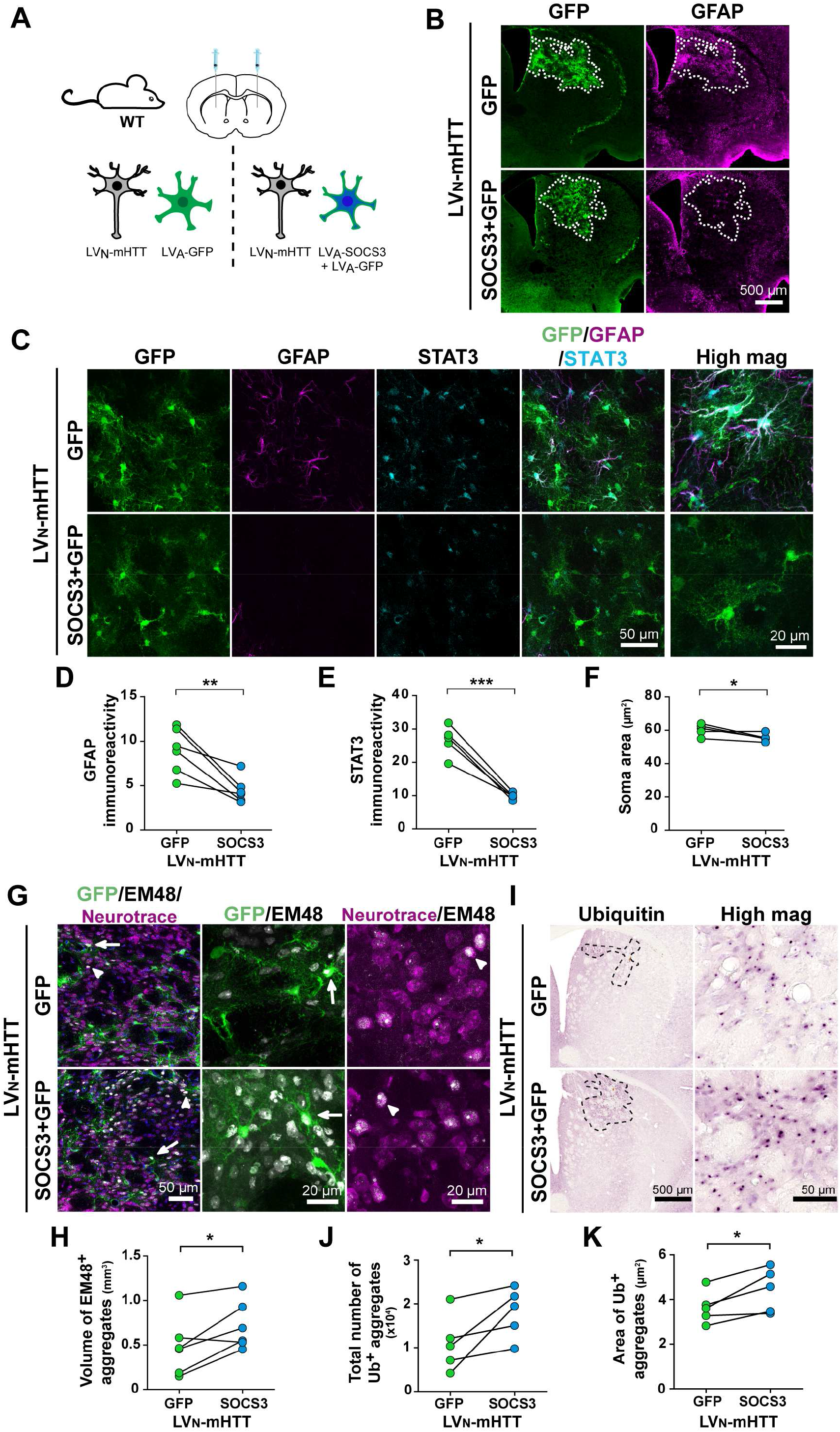
SOCS3-inhibition of the JAK2-STAT3 pathway in astrocytes increases the number and size of neuronal mHTT aggregates A. Two month-old WT mice were injected with LVN-mHTT + LVA-GFP in one striatum and with LVN-mHTT + LVA-SOCS3 + LVA-GFP in the contralateral striatum, at the same total virus load. Their brains were analysed 6 weeks later. **B.** Low magnification images show the transduced GFP^+^ area (green, outlined) in the striatum and immunostaining for GFAP (magenta). **C.** High magnification confocal images of astrocytes stained for GFP (green), GFAP (magenta) and STAT3 (cyan). **D-F**. Immunoreactivity for GFAP (**D**) and STAT3 (**E**), as well as astrocyte soma area (**F**) are significantly decreased by SOCS3. Paired *t*-test. *n* = 5-6/group, **D**: *df* = 5, *t* = 4.349, *p* = 0.0074; **E**: *df* = 4, *t* = 8.651, *p* = 0.0010; **F**: *df* = 4, *t* = 2.961, *p* = 0.0415. **G**. The striatal volume with EM48^+^ aggregates is significantly increased by SOCS3. Paired *t*- test. *n* = 6/group, *df* = 5, *t* = 3.096, *p* = 0.0270. **H.** Confocal images of striatal sections stained for GFP (green), EM48 (white), neurotrace (magenta) and DAPI (blue). Large EM48^+^ aggregates of mHTT are mostly found in neurons stained for neurotrace, occupying their entire nucleus (arrowhead). Only few GFP^+^ astrocytes display an EM48^+^ aggregate (arrow). **I.** Immunolabeling for ubiquitin (Ub) shows Ub^+^ aggregates in the striatum (delimited with black dots). **J.** The number of Ub^+^ inclusions is significantly increased by SOCS3. Wilcoxon matched-pair test, *n* = 5/group, *p* = 0.0431. **K.** Ub^+^ inclusions are larger in the striatum injected with LVA-SOCS3 than LVA-GFP. Paired *t*-test. *n* = 5/group, *df* = 4, *t* = 3.406, *p* = 0.0271.

### Activation of JAK2-STAT3 pathway in astrocytes is beneficial for striatal neurons

We next evaluated how JAK2-STAT3 pathway activation in astrocytes impacted several disease outcomes. Hdh140 mice do not show the typical striatal neurodegeneration observed in Huntington’s disease patients, but display early transcriptional defects in striatal neurons, in particular for *Ppp1r1b* transcripts (which encode the striatal protein DARPP32). In Hdh140 mice, JAK2ca expression in striatal astrocytes significantly increased levels of the neuronal transcript *Ppp1r1b* (**Fig. 4A**), suggesting a beneficial effect on neurons. To further demonstrate JAK2ca beneficial effects on clinically-relevant pathological outcomes, we next performed magnetic resonance imaging. Hdh140 mice display striatal atrophy, which is detectable with T2-weighted magnetic resonance imaging^46^, and is observed at prodromal stages in Huntington’s disease patients^47^. Hdh140 mice also display lower striatal levels of the major neurotransmitter and metabolic intermediate glutamate^46^, again replicating an early pathological hallmark of Huntington’s disease patients^48^. Brain glutamate distribution can be mapped with a high resolution using glutamate Chemical Exchange Saturation Transfer (gluCEST)^49^, revealing local reduction in glutamate abundance in Hdh140 mice^50^. Hdh140 mice were injected in one striatum with LVA-JAK2ca + LVA-GFP and the other striatum with LVA-GFP as an internal control, and imaged 2 months later. The volume of the striatum injected with LVA-JAK2ca was significantly larger than the contralateral striatum injected with LVA- GFP (**Fig. 4B**), showing reduced atrophy with JAK2ca. Likewise, gluCEST imaging revealed significantly higher striatal levels of glutamate with JAK2ca (**Fig. 4C**), further supporting beneficial effects of JAK2-mediated astrocyte reactivity on two pathological hallmarks of Huntington’s disease.

**Figure 4.**
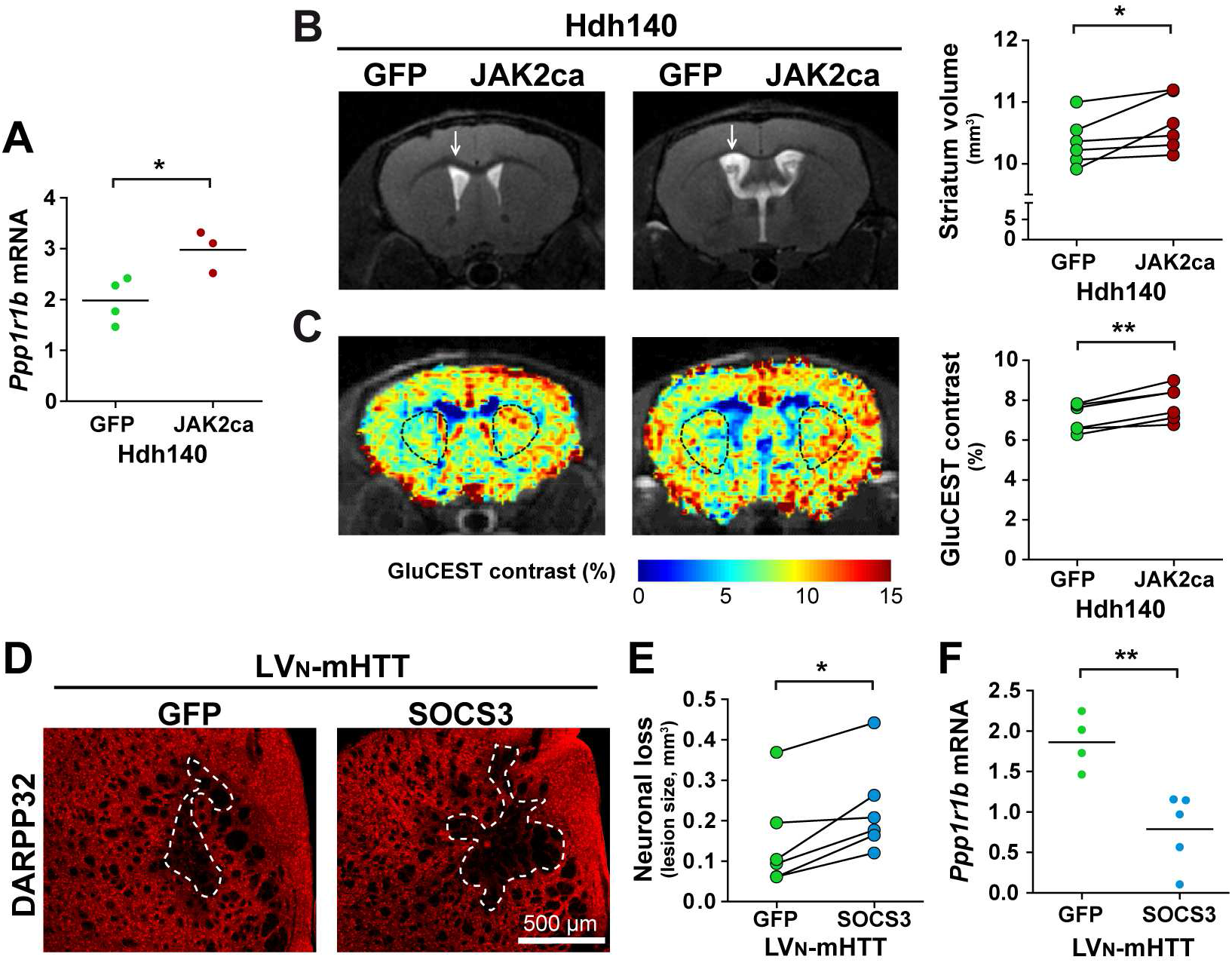
Activation of the JAK2-STAT3 pathway in astrocytes has beneficial effects on striatal neurons A. Mice were injected as in Fig. 2A. Striatal *Ppp1r1b* mRNA levels are higher in Hdh140- JAK2ca mice than in Hdh140-GFP mice. Student *t*-test. *n* = 3-5/group, *df* = 5, *t* = 3.028, *p* = 0.0291. **B.** T2-weigthed magnetic resonance images show that LVA-JAK2ca reduces striatal atrophy in Hdh140 mice. Note the larger ventricle in the side injected with the control vector LVA-GFP, caused by striatal atrophy (arrow). Wilcoxon matched-pair test, *p* = 0.0313. **C.** Quantification of gluCEST contrast in Hdh140 mice reveals higher glutamate levels in the striatum injected with LVA-JAK2ca. Paired *t*-test, *n* = 6, *df* = 5, *t* = 5.641, *p* = 0.0024. Two representative mice are shown, each on a different anatomical position. Region of interest are shown with black dash on each striatum. **D-F** Mice were injected as in Fig. 3A. **D.** Images of striatal sections stained for DARPP32 (red) showing the striatal lesion caused by mHTT (white dotted lines). **E.** Striatal DARPP32^-^ lesions are significantly larger in the SOCS3 group. Wilcoxon matched-pair test, *n* = 6/group, *p* = 0.0277. **F.** SOCS3 decreases mRNA levels of the neuronal transcripts *Ppp1r1b* (*Darpp32).* Student *t*-test. *n* = 4-5/group, *df* = 7, *t* = 3.931, *p* = 0.0057.

In the lentiviral model, mHTT causes local neuronal degeneration, visible as DARPP32- depleted lesion (**Fig. 4D**). SOCS3 significantly increased the lesion volume (**Fig. 4E**), and reduced *Ppp1r1b* mRNA levels (**Fig. 4F**).

Overall, our data show that activation of the JAK2-STAT3 pathway in reactive astrocytes reduces the number and size of mHTT aggregates in neurons and mitigates Huntington’s disease neuronal alterations; while blocking the pathway has opposite effects.

### JAK2ca regulates the expression of proteostasis genes in astrocytes

How can JAK2-STAT3 pathway activation in reactive astrocytes impact mHTT aggregation in neurons? As the JAK2-STAT3 cascade regulates gene expression, we investigated transcriptional changes induced by JAK2ca by comparing the transcriptome of acutely sorted astrocytes isolated from WT-JAK2ca and WT-GFP control mice by microarray (**Fig. 5A**). Transduced astrocytes were collected by fluorescence-activated cell sorting (FACS) based on their GFP expression. GFP^-^ cells, comprising microglia, neurons, oligodendrocyte precursor cells (OPC), oligodendrocytes and few non-infected astrocytes, were collected together. There were 1,415 differentially expressed transcripts (fold change > 1.5, *p* < 0.05), between GFP^+^ and GFP^-^ cell samples in control WT-GFP mice (**Fig. 5B**). Besides *eGfp*, many known astrocyte gene markers were enriched in GFP^+^ cells (e.g. *AldoC, Aqp4, Gja1, Gjb6, Slc1a2, Slc1a3*). Conversely, known markers for microglial cells, neurons, OPC, oligodendrocytes were enriched in GFP^-^ cells (e.g. *Aif, Trem2, P2ry12, Pde10a, Snap25, Pdgfra, Myt1*). The sorting procedure being validated, we next compared the gene expression profile of GFP^+^ astrocytes isolated from WT-GFP and WT-JAK2ca mice.

**Figure 5.**
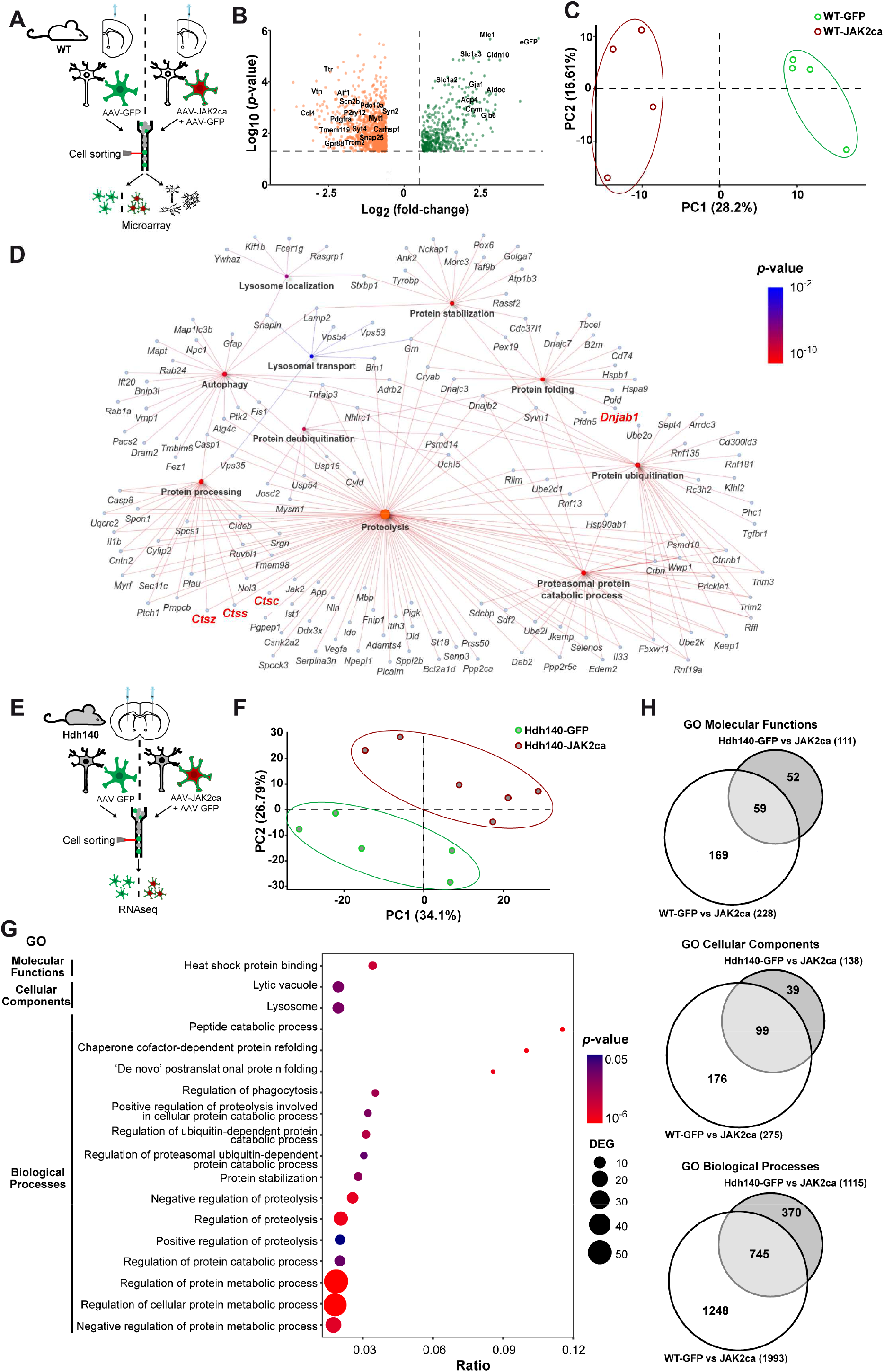
JAK2ca regulates the expression of proteostasis genes in astrocytes A. Two month-old WT mice were injected in the striatum with AAV-GFP or AAV-JAK2ca + AAV-GFP (N = 4/group), at the same total virus load. After 2 months, GFP^+^ striatal astrocytes were acutely sorted and their transcriptome analysed by microarray. **B.** Validation of astrocyte isolation. The volcano plot shows the 1,415 differentially expressed genes between GFP^+^ astrocytes and GFP^-^ cells in WT-GFP mice (in green overexpressed in GFP^+^ cells; in orange, overexpressed in GFP^-^ cells). eGFP and established cell-type specific markers are shown. GFP^+^ cells express typical astrocyte markers while GFP^-^ cells express markers for microglial cells, neurons, cells of the oligodendrocyte lineage and endothelial cells. **C.** Principal component analysis shows a clear segregation of the four WT-GFP and four WT-JAK2ca astrocyte samples. **D**. GO analysis on the list of JAK2ca-regulated genes in GFP^+^ astrocytes reveals a significant enrichment in several biological processes linked to autophagy/lysosome or UPS and in molecular functions linked to chaperones. Network plot with the genes involved in the selected GO pathways. *n* = 4/group. **E.** Hdh140 mice (8-10 month-old) were injected in the striatum with AAV-GFP or AAV-JAK2ca + AAV-GFP, at the same total virus load. After 4 months, GFP^+^ striatal astrocytes were collected and analysed by RNAseq. **F.** The five Hdh140- GFP samples cluster apart from the six Hdh140-JAK2ca astrocyte samples, on a principal component analysis graph. **G.** GO analysis reveals a significant enrichment in genes involved in molecular functions (MF), cellular components (CC) and biological processes (BP) linked to proteostasis. The x axis represents the ratio of the number of differentially expressed genes over the total number of genes belonging to a GO entry, DEG = differentially expressed genes. *n* = 5-6/group. **H**. Venn diagrams for the enriched GO molecular functions, cellular components and biological functions, from the list of differentially expressed genes in WT-JAK2ca and Hdh140-JAK2ca astrocytes. There is a majority of common terms.

Astrocyte samples from these two groups clustered separately by principal component analysis (**Fig. 5C**). We found 888 probes (802 unique transcripts) differentially expressed between JAK2ca-astrocytes and control GFP-astrocytes, including *Jak2* mRNA itself (**Supplementary Fig. 2F**). A Gene Ontology (GO) analysis revealed a significant enrichment in many GO-biological processes linked to immunity and inflammation, confirming that JAK2ca triggers reactive changes in astrocytes, which were also evidenced by morphological changes (**Supplementary Fig. 2A-H**)^39^.

Among the differentially expressed genes between GFP- and JAK2ca-astrocytes, there was a specific enrichment in biological processes linked to lysosomes and UPS, as well as other processes related to proteostasis (**Fig. 5D**). KEGG pathway analysis also revealed a significant enrichment in the term “lysosome” in JAK2ca-reactive astrocytes (*p* = 0.0005). Several cathepsins (*Ctsc, Ctss* and *Ctsz*) were upregulated by JAK2ca in astrocytes (**Fig. 5D**). Genes linked to proteostasis formed a complex network of co-regulated genes in JAK2ca-astrocytes (**Fig. 5D**).

To confirm that the JAK2-STAT3 pathway was also able to induce a proteostasis gene signature in astrocytes in a Huntington’s disease context, we sorted striatal astrocytes from Hdh140-GFP and Hdh140-JAK2ca mice based on their GFP expression, and performed RNAseq analysis (**Fig. 5E**). Again, sorted astrocytes expressed high levels of astrocyte-specific genes and low or undetectable levels of known markers for microglia, neurons, cells of the oligodendrocyte lineage and endothelial cells (**Supplementary Fig. 2I**). Principal component analysis showed that astrocytes from Hdh140-GFP and Hdh140-JAK2ca mice formed two distinct clusters (**Fig. 5F**). We found 269 genes differentially expressed between Hdh140-GFP and Hdh140-JAK2ca astrocytes. *Jak2* levels were significantly higher in JAK2ca-astrocytes than GFP-astrocytes (**Supplementary Fig. 2J)**. Murine *Htt,* on the contrary, was expressed at low levels in sorted astrocytes from both groups (rpkm value = 1.859 and 2.120 for GFP- and JAK2ca-astrocytes; adjusted *p*-value = 0.999), showing that JAK2ca does not change *Htt* transcription in striatal astrocytes. Among the 269 differentially expressed genes in Hdh140-JAK2ca astrocytes, 212 genes were present in the full list of 30,845 unique probes detected by microarray analysis of WT striatal astrocytes (i.e. striatal astrocyte transcriptome). And among these 212 genes, only 20 (10%) were also differentially expressed in WT-JAK2ca astrocytes compared to WT-GFP astrocytes. However there was a significant convergence on the functions of the differentially expressed genes between the two studies. In particular, many of the differentially expressed genes in Hdh-140-JAK2ca astrocytes were linked to immunity/inflammation, as found in WT- JAK2ca astrocytes (**Supplementary Fig. 2E**). Likewise, as in WT-JAK2ca astrocytes, there was a significant enrichment in GO pathways related to proteostasis, including the molecular function “Heat shock protein binding” and the cellular components “lytic vacuole” and “lysosome” (**Fig. 5G**). Gene Set Enrichment Annotation (GSEA) also identified the term “phagosome”, as significantly enriched in Hdh140-JAK2ca astrocytes, with a majority of up- regulated genes (normalized enrichment score = 1.759, adjusted *p* value = 0.039). Globally, we found that 53% of GO molecular functions, 71% of GO cellular components and 67% of GO biological processes were shared between Hdh140-JAK2ca astrocytes and WT-JAK2ca astrocytes (**Fig. 5H**).

Overall, this transcriptomic analysis shows that JAK2ca induces a specific proteostasis gene signature in striatal astrocytes both in WT and Hdh140 mice.

### JAK2ca increases proteolytic capacity in Huntington’s disease astrocytes

It is important to establish that the identified transcriptional changes translate into detectable changes in astrocyte function^51^. To assess proteolytic activity of the two major clearance pathways specifically in astrocytes of Huntington’s disease mice, we used cell-permeable, activity probes for the lysosomal enzymes cathepsins and for the proteasome (**Supplementary Fig. 1B**).

We incubated acutely dissociated striatal cells from Hdh140-GFP and Hdh140-JAK2ca mice in a pan-cathepsin activity probe that becomes fluorescent when metabolized by cathepsins^52^. We then measured probe fluorescence in GFP^+^ astrocytes from the two groups by FACS (**Fig. 6A)**. There was a significantly larger fraction of GFP^+^ astrocytes with high cathepsin activity in Hdh140-JAK2ca mice (**Fig. 6B**), revealing that JAK2ca increases lysosomal activity in astrocytes.

**Figure 6.**
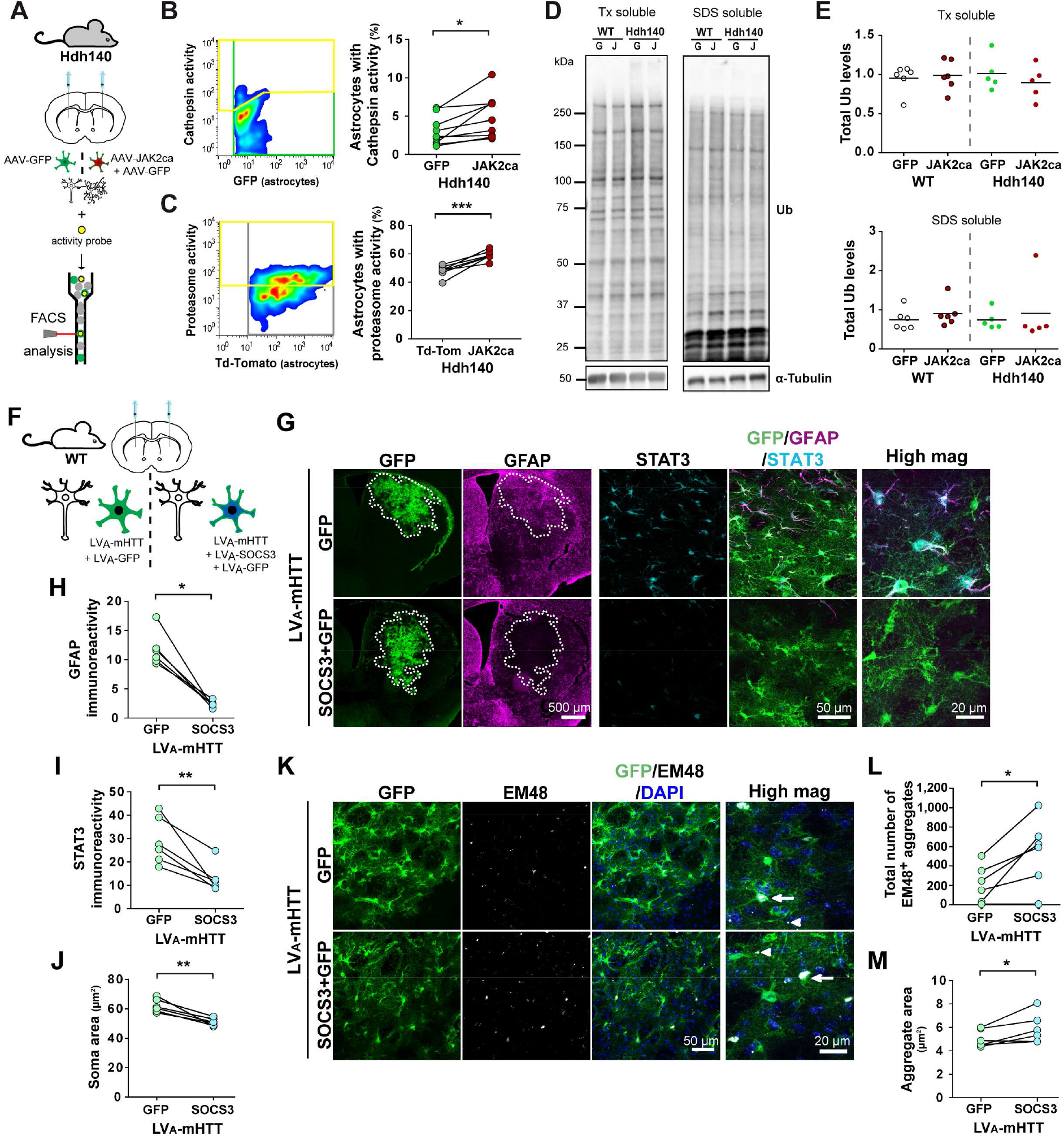
JAK2ca increases cathepsin and proteasome intrinsic activities in Huntington’s disease striatal astrocytes A-C. The striatum of Hdh-JAK2ca mice and their control Hdh140-GFP or Hdh140-Td-Tomato was collected, cells dissociated and incubated with a quenched fluorescent pan-cathepsin activity probe (**B**) or a proteasome activity probe (**C**). The percentage of GFP^+^ astrocytes with cathepsin activity (**B**) or the percentage of Td-Tomato^+^ astrocytes with proteasome activity (**C**) was quantified in each mouse. Hdh140-JAK2ca mice display a higher percentage of astrocytes with cathepsin or proteasome activity. Paired *t*-test on *arcsine-*transformed data. **C**: *n* = 9/group, *df* = 8, *t* = 3.183, *p* = 0.0129. **D**: *n* = 7/group, *df* = 6, *t* = 6.001, *p* = 0.0010. **D, E.** Immunoblotting for Ub was performed on striatal Tx-soluble and SDS-soluble fractions from WT-GFP, WT- JAK2ca, Hdh140-GFP and Hdh140-JAK2ca mice. Immunoreactivity pattern and total Ub levels are not different between groups. Band intensity was normalized to α-tubulin. For Tx- soluble: Two-way (genotype, treatment) ANOVA, *n* = 5-6/group, *df* = 1, Genotype: F = 0.038, *p* = 0.8481; Treatment: F = 0.224, *p* = 0.6421, Genotype x Treatment: F = 0.776, *p* = 0.3900. For SDS-soluble: Kruskal-Wallis test *n* = 5-6/group, *p* = 0.3211. **F-M.** Two month-old WT mice were injected in one striatum with LVA-mHTT and LVA-GFP and the contralateral striatum with LVA-mHTT, LVA-SOCS3 and LVA-GFP (same total viral load) to force mHTT expression in astrocytes. Their brains were analysed 6 weeks later. **G.** Low magnification images showing GFP^+^ transduced area (outlined, green) and GFAP staining (magenta) in the mouse striatum. Confocal images of striatal sections stained for GFP (green), GFAP (magenta) and STAT3 (cyan). SOCS3 reduces GFAP immunoreactivity and nuclear accumulation of STAT3 in astrocytes. **H-J**. Quantification of GFAP immunoreactivity (**H**), STAT3 immunoreactivity (**I**) and astrocyte soma area (**J**). SOCS3 significantly decreases all these parameters. **H**: Wilcoxon matched-pair test, *n* = 6/group, *p* = 0.0277. **I**: Paired *t*-test. *n* = 6/group, *df* = 5, *t* = 4.354, *p* = 0.0073. **J**: Paired *t*-test. *n* = 6/group, *df* = 5, *t* = 5.442, *p* = 0.0028. **K**. Confocal images of striatal sections stained for GFP (green), EM48 (white) and DAPI (blue). Large EM48^+^ aggregates form in astrocyte nucleus (arrow), while small aggregates are mainly found in astrocyte processes (arrowhead). **L, M**. The total number (**L**) and the size (**M**) of EM48^+^ aggregates are significantly increased by SOCS3. Paired *t*-test. *n* = 6/group, *df* = 5, **L**: *t* = 3.246, *p* = 0.0228. **M**: *t* = 2.778, *p* = 0.0390.

Another fluorescent probe was used to measure proteasome activity in acutely dissociated astrocytes (**Fig. 6C**)^53^. As the excitation/emission spectrum of this probe overlaps with GFP, we used a viral vector encoding the red fluorescent protein Td-Tomato instead of GFP to detect astrocytes in both groups. Again, the fraction of Td-Tomato^+^ astrocytes with high proteasome activity was larger in Hdh140-JAK2ca mice than in Hdh140-Td-Tomato mice (**Fig. 6C**), showing that reactive astrocytes also have a higher proteasome activity.

As several ubiquitin ligases were differentially expressed in JAK2ca-astrocytes (**Fig. 5D**), we assessed ubiquitination by immunoblotting striatal homogenates with a Ub antibody. We did not observe major changes in the pattern of Ub immunoreactivity or in total Ub levels between Hdh140-GFP and Hdh140-JAK2ca mice, both in Tx- and SDS-soluble fractions (**Fig. 6D, E**), suggesting that only the proteolytic step of the UPS is stimulated by JAK2ca, without global changes in the ubiquitination profile of mouse striatum.

To directly measure the intrinsic capacity of astrocytes to clear mHTT, we used viral vectors to force mHTT expression in astrocytes (LVA-mHTT). WT mice were injected in the right striatum with LVA-mHTT, LVA-SOCS3, and LVA-GFP, and in the control left striatum with LVA-mHTT and LVA-GFP, at the same total viral titer (**Fig. 6F**). Expression of mHTT in astrocytes also triggered STAT3 activation, as evidenced by its nuclear accumulation, and induced reactive changes in astrocytes (**Fig. 6G**)^8^. In this model as well, SOCS3 efficiently reduced GFAP levels (**Fig. 6G, H**), STAT3 nuclear accumulation (**Fig. 6G, I)** and astrocyte soma hypertrophy (**Fig. 6G, J**). If the JAK2-STAT3 pathway enhances proteolytic activity in astrocytes, we reasoned that blocking this pathway with SOCS3 would increase mHTT aggregation in astrocytes. Indeed, we observed that the total number of mHTT aggregates in the striatum was increased with SOCS3 (**Fig. 6K, L**). Moreover, SOCS3 increased mHTT aggregate size (**Fig. 6K, M**). In this model, mHTT forms both nuclear and cytoplasmic inclusions in astrocytes (**Fig. 6K**). The fraction of nuclear aggregates in GFP^+^ astrocytes was not changed by SOCS3 (19.1% and 22.0% respectively in GFP and SOCS3 groups, Student *t-* test on *arcsine*-transformed data, *n* = 6/group, *df* = 5, *t* = 0.787*, p* = 0.458). These results show that the JAK2-STAT3 pathway stimulates astrocyte intrinsic capacity for mHTT clearance.

### JAK2ca induces chaperone expression in astrocytes

Interestingly, several GO-molecular functions linked to chaperones and protein folding were significantly regulated by JAK2ca both in WT and Hdh140 astrocytes (**Fig. 5D, G**). Chaperones prevent mHTT aggregation^32, 54^ and can be released extracellularly in exosomes^55^.

We focused on the co-chaperone DNAJB1 [DnaJ heat shock protein family (Hsp40) member B1], a member of the HSP40 family, which is induced nearly 3-fold by JAK2ca in WT astrocytes (**Fig. 5D**). DNAJB1 immunoreactivity was higher in the putamen of Huntington’s disease patients, specifically at the core of the degenerative area devoid of NeuN^+^ neurons and with abundant hypertrophic GFAP^+^ astrocytes (**Fig. 7A**).

**Figure 7.**
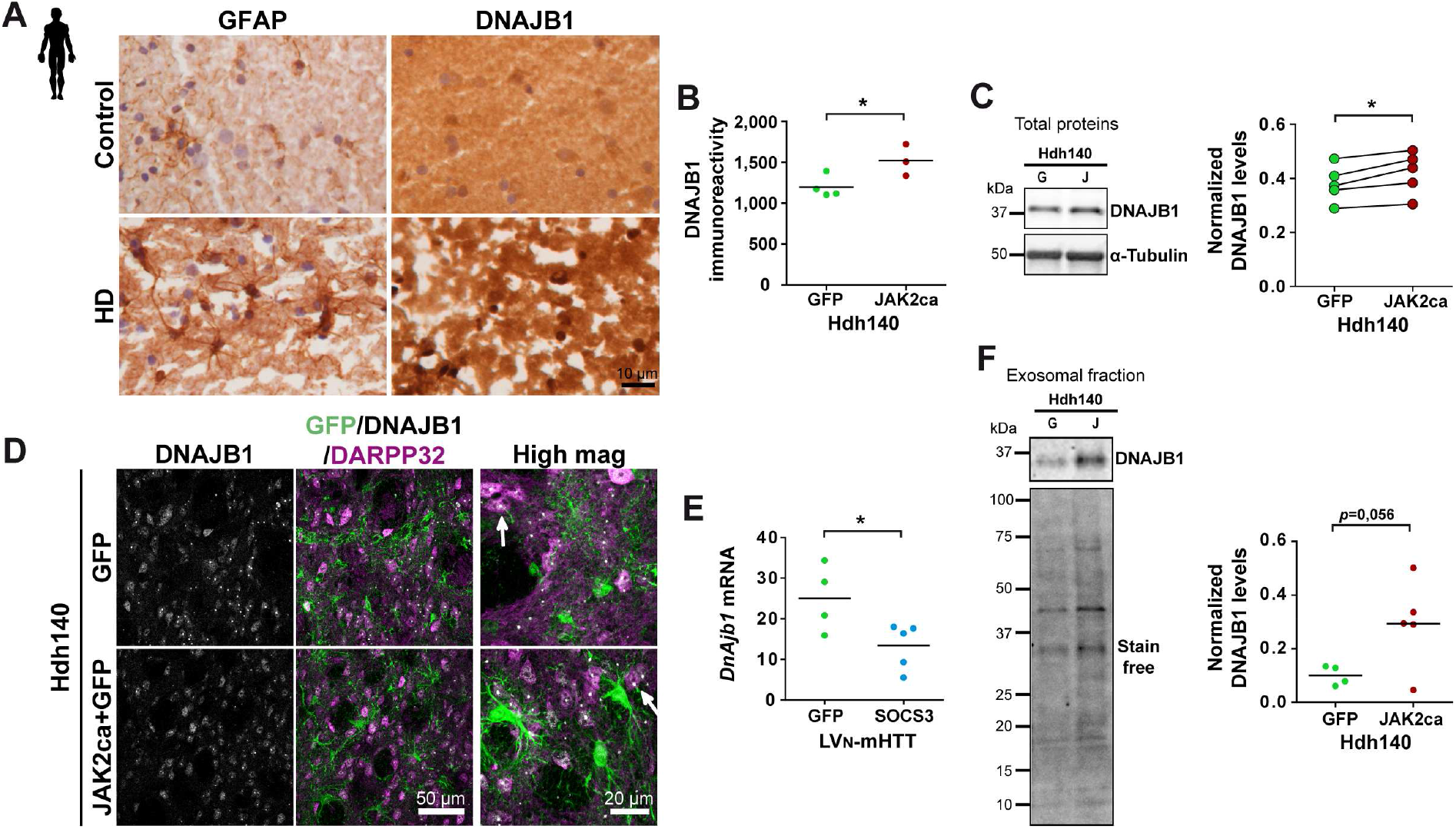
The JAK2-STAT3 pathway increases DNAJB1 expression in astrocytes A. DNAJB1 display a strong immunoreactivity in the degenerative putamen region that is filled with GFAP^+^ astrocytes in Huntington’s disease patients. **B, C.** Immunostaining (**B**) and immunoblotting (**C**) show higher DNAJB1 protein levels in the striatum of Hdh140-JAK2ca mice than Hdh140-GFP mice. Band intensity was normalized to α-tubulin. **B**: Student *t*-test. *n* = 3-5/group, *df* = 5, *t* = 2.658, *p* = 0.0450. **C**: Paired *t*-test. *n* = 5/group, *df* = 4, *t* = 4.079, *p* = 0.0151. **D**. Confocal images of striatal sections stained for GFP (green), DNAJB1 (white) and DARPP32 (magenta). DNAJB1 displays a diffuse cytosolic staining and forms small nuclear inclusions in neurons (arrow) of Hdh140 mice. **E.** *Dnajb1* mRNA levels are significantly decreased by SOCS3 in LVN-mHTT mice. Student *t*-test. *n* = 4-5/group, *df* = 7, *t* = 2.525, *p* = 0.0395. **F.** DNAJB1 is present in exosomes isolated from Hdh140 striata and its levels, normalized by stain free staining, tend to be higher in Hdh140-JAK2ca than Hdh140-GFP exosomes. Student *t*-test. *n* = 4-5/group, *df* = 7, *t* = 2.290, *p* = 0.0558.

JAK2ca significantly increased DNAJB1 protein levels in the striatum of Hdh140 mice as seen by immunostaining (**Fig. 7B, D**) and immunoblotting (**Fig. 7C**), while *Dnajb1* mRNA levels were significantly reduced by SOCS3 in the lentiviral model of Huntington’s disease (**Fig. 7E**). In Hdh140 mice, DNAJB1 displayed a diffuse cytosolic staining but also formed small nuclear inclusion-like structures, suggesting that DNJAB1 can be in close association with mHTT aggregates (**Fig. 7D**).

We next studied whether DNAJB1 was found in exosomes and whether DNAJB1 exosomal content was impacted by JAK2-STAT3 activation in Hdh140 mice. Exosomal vesicles were isolated by biochemical fractionation from the striatum of Hdh140-GFP and Hdh140-JAK2ca mice, and they did contain DNAJB1 (**Supplementary Fig. 1C**). DNAJB1 normalized levels displayed a strong tendency to be higher in exosomes of Hdh140-JAK2ca mice than in Hdh140- GFP control mice (*p* = 0.056, **Fig. 7F**).

To assess whether DNAJB1 released by JAK2ca-astrocytes contributes to reduce mHTT aggregation in neurons, we generated viral vectors targeting astrocytes and encoding a dominant-negative form of human *DNAJB1* (DNAJB1-DN), which prevents DNAJB1 interaction with the HSP70 chaperone^42^, without impacting its loading into exosomes^55^. DNAJB1-DN was expressed in astrocytes in Hdh140-JAK2ca and Hdh140-GFP mice (**Fig. 8A**). DNAJB1-DN, detected by its V5 tag, was confirmed to be expressed in striatal GFP^+^ astrocytes, throughout their cell body and fine processes (**Fig. 8B**). Interestingly, small V5^+^ cytoplasmic vesicles were also observed (although rarely) in nearby DARPP32^+^ neurons (**Fig. 8C**). Some vesicles were co-labelled with the endo-lysosomal marker LAMP1 (**Supplementary Fig. 1D**), suggesting that DNAJB1 can indeed be shuttled from astrocytes to neurons through exosomes. JAK2ca was still able to increase GFAP levels in Hdh140 astrocytes in presence of DNAJB1-DN (**Fig. 8D**). However, co-expression of DNAJB1-DN blocked JAK2ca-mediated reduction of EM48^+^ aggregate numbers (**Fig. 8E**) and even decreased *Ppp1r1b* mRNA levels (**Fig. 8F**).

**Figure 8.**
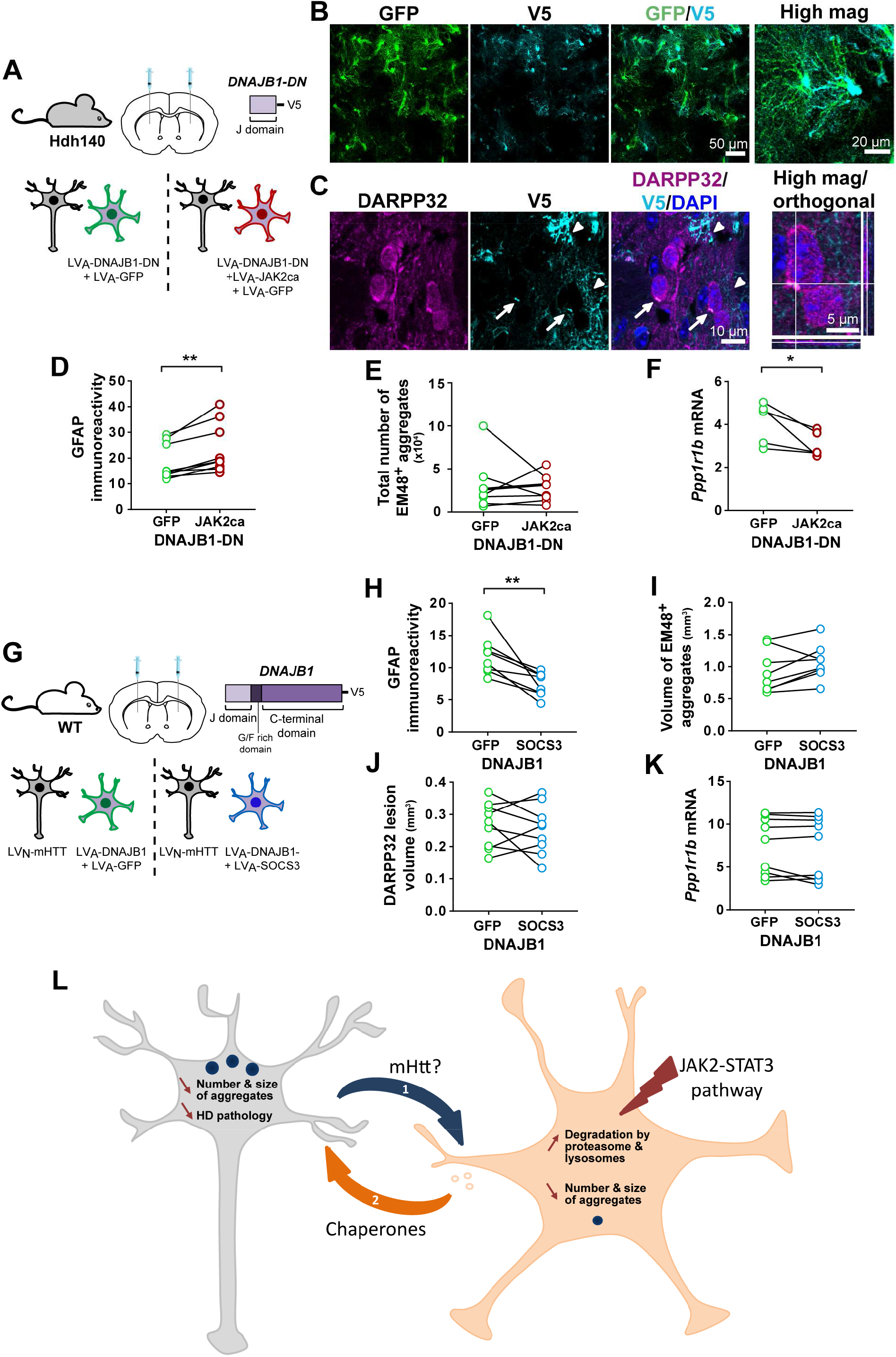
DNAJB1 is involved in the anti-aggregation effects of the JAK2-STAT3 pathway A. Hdh140 mice (8-9 month-old) were injected in one striatum with LVA-GFP + LVA- DNAJB1-DN and with LVA-JAK2ca + LVA-GFP + LVA-DNAJB1-DN in the contralateral striatum, at the same total virus load. Their brains were analysed 4 months later. **B.** Representative confocal images showing striatal sections stained for GFP (green) and V5 (cyan) in astrocytes. **C**. V5^+^ vesicles (cyan, arrows) are observed in the cytoplasm of DARPP32^+^ striatal neurons (magenta). High magnification of a single z plane and orthogonal views are shown on the right, DAPI is shown in blue. Contrary to infected astrocytes expressing high levels of V5 throughout their cytoplasm, including in their fine processes infiltrating the neuropil (arrowheads), neurons display V5 staining only in few cytoplasmic vesicles, ruling out a direct infection of striatal neurons by LVA-DNAJB1-DN. Additional negative controls are shown in **Supplementary** Fig. 1D**. D.** JAK2ca increases GFAP immunoreactivity in astrocytes overexpressing DNAJB1-DN in Hdh140 mice. Wilcoxon matched-pair test, *n* = 8/group, *p* = 0.0077. **E.** In presence of DNAJB1-DN, the total number of EM48^+^ aggregates is no longer decreased by JAK2ca. Wilcoxon matched-pair test, *n* = 8/group, *p* = 0.7794. **F.** *Ppp1r1b* mRNA levels are significantly reduced by JAK2ca in Hdh140 mice expressing DNAJB1-DN. Wilcoxon matched-pair test, *n* = 8/group, *p* = 0.0431. **G.** Two month-old WT mice were injected in one striatum with LVN-mHTT + LVA-GFP + LVA-DNAJB1 and the contralateral striatum with LVN-mHTT + LVA-SOCS3 + LVA-DNAJB1, at the same total virus load and analysed 6 weeks later. **H.** SOCS3 reduces GFAP immunoreactivity in LVN-mHTT mice, even in presence of DNJAB1. Paired *t*-test. *n* = 9/group, *df* = 8, *t* = 3.534, *p* = 0.0077. **I**. SOCS3 no longer increases the EM48^+^ volume when astrocytes co-express DNAJB1. Paired *t*-test. *n* = 9/group, *df* = 8, *t* = 2.004, *p* = 0.0851. **J, K.** Likewise, striatal DARPP32^-^ lesions (**J**) and *Ppp1r1b* mRNA levels (**K**) are no longer different between the two groups. **J**: Paired *t*-test. *n* = 9/group, *df* = 8, *t* = 0.632, *p* = 0.5451. **K:** Wilcoxon matched-pair test, *n* = 8/group, *p* = 0.8203. **L. Bi- directional communication between JAK2-STAT3-induced reactive astrocytes and neurons to promote proteostasis in Huntington’s disease:** The JAK2-STAT3 pathway induces a reactive response in striatal astrocytes and activates a transcriptional program that promotes proteostasis, reduces mHTT aggregation and improves neuronal status. Two complementary and non-exclusive mechanisms involving striatal neurons and JAK2-STAT3- dependent reactive astrocytes may take place. **1.** Reactive astrocytes display a higher intrinsic proteolytic activity that promotes mHTT degradation. Neurons could shuttle their mHTT to astrocytes for clearance. The exact mechanisms and the form of mHTT exchanged (monomers, fibrils, cleaved fragments…) remain to be determined. **2.** Reactive astrocytes over-express chaperones such as DNAJB1 that can be released in exosomes towards neurons to promote their proteostasis.

Conversely, we tested whether DNAJB1 restoration was able to oppose SOCS3 detrimental effects in the lentiviral model of mHTT overexpression. In the striatum of WT mice, we injected a viral vector targeting astrocytes and encoding a full length human *DNAJB1* (LVA-DNAJB1) with LVA-SOCS3 or LVA-GFP together with LVN-mHTT (**Fig. 8G**). DNAJB1 did not interfere with SOCS3-mediated reduction of GFAP levels in Huntington’s disease astrocytes (**Fig. 8H**). However, when DNAJB1 was expressed in astrocytes, SOCS3 no longer exacerbated mHTT aggregation (**Fig. 8I**), neuronal lesion (**Fig. 8J**), or reduced *Ppp1r1b* mRNA levels (**Fig. 8K**), suggesting that DNAJB1 expression in astrocytes counteracts SOCS3 deleterious effects.

Our results suggest that the co-chaperone DNAJB1 produced by JAK2-STAT3 reactive astrocytes helps reduce mHTT aggregation and neuronal alterations in Huntington’s disease.

## Discussion

We studied how the JAK2-STAT3 pathway shapes the proteostasis response of reactive astrocytes in Huntington’s disease. We found that STAT3 is activated in reactive astrocytes of Huntington’s disease patients. Thanks to two complementary mouse models of Huntington’s disease allowing the selective and reciprocal manipulation of the JAK2-STAT3 pathway in striatal astrocytes, we show that this cascade controls astrocyte reactive state and reduces both the number and size of mHTT aggregates forming in neurons.

The reduction of mHTT aggregation by JAK2-STAT3 pathway activation in astrocytes is not due to lower mHTT expression, as *Htt* mRNA levels and Tx-soluble mHTT concentrations were not reduced by JAK2ca. Importantly, JAK2ca-mediated reduction in mHTT aggregation did not trigger an increase in soluble mHTT levels either. This observation rules out that JAK2ca prevents mHTT from coalescing into aggregates or favours the accumulation of soluble mHTT after its dissociation from aggregates. Instead, activation of the JAK2-STAT3 pathway in reactive astrocytes appears to promote the full degradation of mHTT insoluble oligomers or aggregates, which could be mediated by autophagy-lysosomal removal of aggregates or chaperone-mediated extraction of mHTT and targeting to the UPS for complete clearance.

In keeping with this, transcriptomic analysis of acutely sorted astrocytes following JAK2- STAT3 pathway activation in WT and Huntington’s disease mice reveals extensive changes in genes linked to lysosomal degradation and the UPS. This result is in accordance with the single nuclei RNAseq analysis of astrocytes from the cingulate cortex of grade III/IV Huntington’s disease patients, reporting a significant enrichment in proteostasis functions^5^. Lysosomes and UPS are active in all brain cells and changes in expression or activity in astrocytes could be masked by larger changes in other cell-types if assessed in typical bulk analyses. We thus implemented two FACS-based assays to measure cathepsin and proteasome activity specifically in astrocytes. We found that JAK2ca increases both proteolytic activities in Huntington’s disease astrocytes. In addition, our ability to induce mHTT expression selectively in astrocytes by viral gene transfer provides a direct demonstration that the JAK2-STAT3 pathway increases reactive astrocyte capacity to clear mHTT.

As aggregates mainly form in neurons, trans-cellular signalling mechanisms must take place between neurons and reactive astrocytes (**Fig. 8L**). Can mHTT be transferred from neurons to reactive astrocytes where they would be degraded more efficiently? In a landmark study in *Drosophila*, it was shown that mHTT exon 1 tagged with mCherry shuttles from neurons to neighbouring phagocytic glia and forms aggregates with endogenous HTT in glia^56^. Other studies showed that mHTT can be exchanged between brain cells in *Drosophila*, in mice^57–59^, and even in humans as mHTT aggregates were detected in healthy embryonic neurons grafted in the brain of Huntington’s disease patients^60^. mHTT can be packaged in exosomes of different cell types^61, 62^ and be taken up by neighbouring cells, including neurons. Exchange of mHTT may also involve direct cell-to-cell contacts via tunnelling nanotubes^63^ or unconventional secretory pathways^64^. Most studies were performed *in vitro* or in non-neuronal cells, therefore, the precise mechanisms of mHTT exchange from neurons to astrocytes in the mammalian brain remain to be elucidated. Of note, a recent study showed that astrocyte-specific silencing of mHTT in a genetic mouse model of Huntington’s disease reduces both astrocyte and neuronal mHTT aggregates^20^, further supporting the concept of a tight partnership between these two cell types to degrade mHTT.

An alternative and non-exclusive mechanism for reduced neuronal mHTT aggregation upon JAK2-STAT3 pathway activation in astrocytes, is that reactive astrocytes release proteins that promote mHTT clearance within neurons (**Fig. 8L**). Our transcriptomic study shows that several chaperones are induced by JAK2ca in reactive astrocytes. In particular, DNAJB1 protein levels were higher in Hdh140-JAK2ca mice and this co-chaperone was abundant in extracellular exosomes, and in the lesioned striatum of Huntington’s disease patients. Chaperones are known to be released in exosomes and mediate trans-cellular proteostasis^34, 55^. Moreover, exosomes isolated from cultured astrocytes were shown to reduce mHTT aggregation in mice^18^. Interestingly, single nuclei RNAseq shows that *DNAJB1*, as well as other chaperones are significantly overexpressed in astrocytes from the cingulate cortex of grade III/IV Huntington’s disease patients^5^ and the putamen of grade II/III patients^9^, showing that this beneficial proteostasis response may also occur in astrocytes from Huntington’s disease patients.

Through its J domain, DNAJB1 interacts with HSP70 to stimulate its ATP-dependent chaperone activity^65^. This domain is also implicated in DNAJB1 loading into exosomes^42, 55^. The J domain alone cannot activate HSP70 and has a dominant-negative action on the endogenous DNAJB1^42^. Expression of this mutant in astrocytes abrogated JAK2ca-mediated beneficial effects in Hdh140 mice, showing DNAJB1 involvement in JAK2ca effects. Conversely, expression of DNAJB1 in astrocytes cancelled SOCS3 deleterious effects on neuronal death and transcriptional defects in the lentiviral model. Small vesicles of tagged DNAJB1 were sometimes observed in neurons nearby infected astrocytes, further supporting the shuttling of DNAJB1 from astrocytes to neurons. Overall, our data strongly suggest that DNAJB1, produced by reactive astrocytes following JAK2-STAT3 pathway activation, promotes mHTT clearance.

DNAJB1 was shown to be the rate-limiting chaperone to suppress aggregation of a short fragment of mHtt^66^. Interestingly, HSPs not only prevent mHTT aggregation but can also solubilize proteins trapped in aggregates like transcription factors and favour mHTT degradation by addressing to the UPS or autophagy-lysosomes^33, 67^. Of note, DNAJB1 was recently shown to promote α-synuclein disaggregation^68^. HSP-mediated extraction of housekeeping proteins or mHTT itself from aggregates is expected to reduce their size, which is consistent with our observations. Therefore, HSPs have multiple actions that can *in fine* protect neurons against mHTT toxicity, as shown in different experimental systems based on HSP overexpression^69–71^. Here, we show that neurons rely on the endogenous production of chaperones by astrocytes to reduce mHTT aggregation.

The toxicity of mHTT aggregates is still discussed^24, 67, 72^ and mHTT aggregates could have a biphasic action^73^. At early stages, they could trap soluble toxic mHTT and prevent its deleterious interaction with key cellular partners. Later, mHTT aggregates could be detrimental by sequestering transcription factors, housekeeping proteins or microRNAs, leading to neuronal dysfunction and necrotic death^24, 74^. It is important to note that most studies were based on *in vitro* systems allowing time-lapse monitoring of aggregates, but which cannot fully replicate the complex brain environment where neurons interact with multiple glial cells and cope with mHTT for months, and even decades in patients. Here, in two complementary mouse models of Huntington’s disease, we report that reduced mHTT aggregation is associated with improved neuronal features, showing that the JAK2-STAT3 pathway shapes a beneficial reactive response in striatal astrocytes. Thanks to sensitive magnetic resonance imaging approaches, we also demonstrate that this pathway reduces striatal atrophy and increases glutamate levels, which are two clinically-relevant disease hallmarks, observed at early stage in patients^47, 48^. Interestingly, we found that the JAK2-STAT3 pathway is still able to activate a protective proteostatic program in Huntington’s disease astrocytes. However, the endogenous activation of the JAK2-STAT3 pathway observed in striatal astrocytes from patients seems to be an insufficient protective response to fully prevent neurodegeneration. Early and widespread stimulation of this astrocytic pathway could enhance neuronal resilience in patients.

In conclusion, we show that the JAK2-STAT3 pathway activates a beneficial proteostasis program in reactive astrocytes, which helps striatal neurons handle toxic mHTT. Our study uncovers two non-mutually exclusive, bi-cellular mechanisms to reduce mHTT aggregation in neurons in Huntington’s disease (**Fig. 8L**). The first mechanism relies on mHTT exit from neurons and clearance within reactive astrocytes and the second involves the release of chaperones from reactive astrocytes to promote neuronal proteostasis. Astrocytes are not only defective in Huntington’s disease as usually reported, they may also acquire enhanced capacities to promote mHTT clearance and neuronal functions, following activation of specific signalling cascades. Our results open new therapeutic avenues to further enhance the natural partnership between reactive astrocytes and vulnerable neurons in Huntington’s disease.

## Acknowledgments

We are grateful to F. Aubry for help with initial AAV vector cloning and FACS experiments, C. Joséphine for AAV production and Dr. J. Baijer and N. Dechamps for FACS-isolation of astrocytes. We thank Prof. Déglon for helpful discussions at the beginning of the project. We acknowledge the help of V. Lavilla, and Cédric Fund on transcriptomic studies at the CNRGH. We thank Drs. K. Cambon and G. Liot for sharing their antibodies to S100β, VDAC and mHTT. We thank L. de Longprez for help with the Hdh140 colony, as well as Dr. N. Heck, E. Saavedra and R. Jacqmin for pilot experiments.

## Competitive interests

The authors declare no competing interests.

## Author contribution

LA, CE: conception; LA, LBH, MACS, CE: design of the work; LA, LBH, PG, MACS, CD, MAP, FP, ASH, MG, MGG, GA, MK, JF: acquisition of data; LA, LBH, MRP, PG, NS, CH, NR, PdlG, JF, RO: analysis of data; LA, LBH, MRP, JF, EBo, SB, RO, EBr, MACS, CE: interpretation of data; MCG, ND, RM, APB, GB: Provided reagents or materials; JFD, PH, EBr, CE: Provided funding; LA, CE: manuscript writing. All authors revised and approved the manuscript.

---

**Supplementary figure 1.**
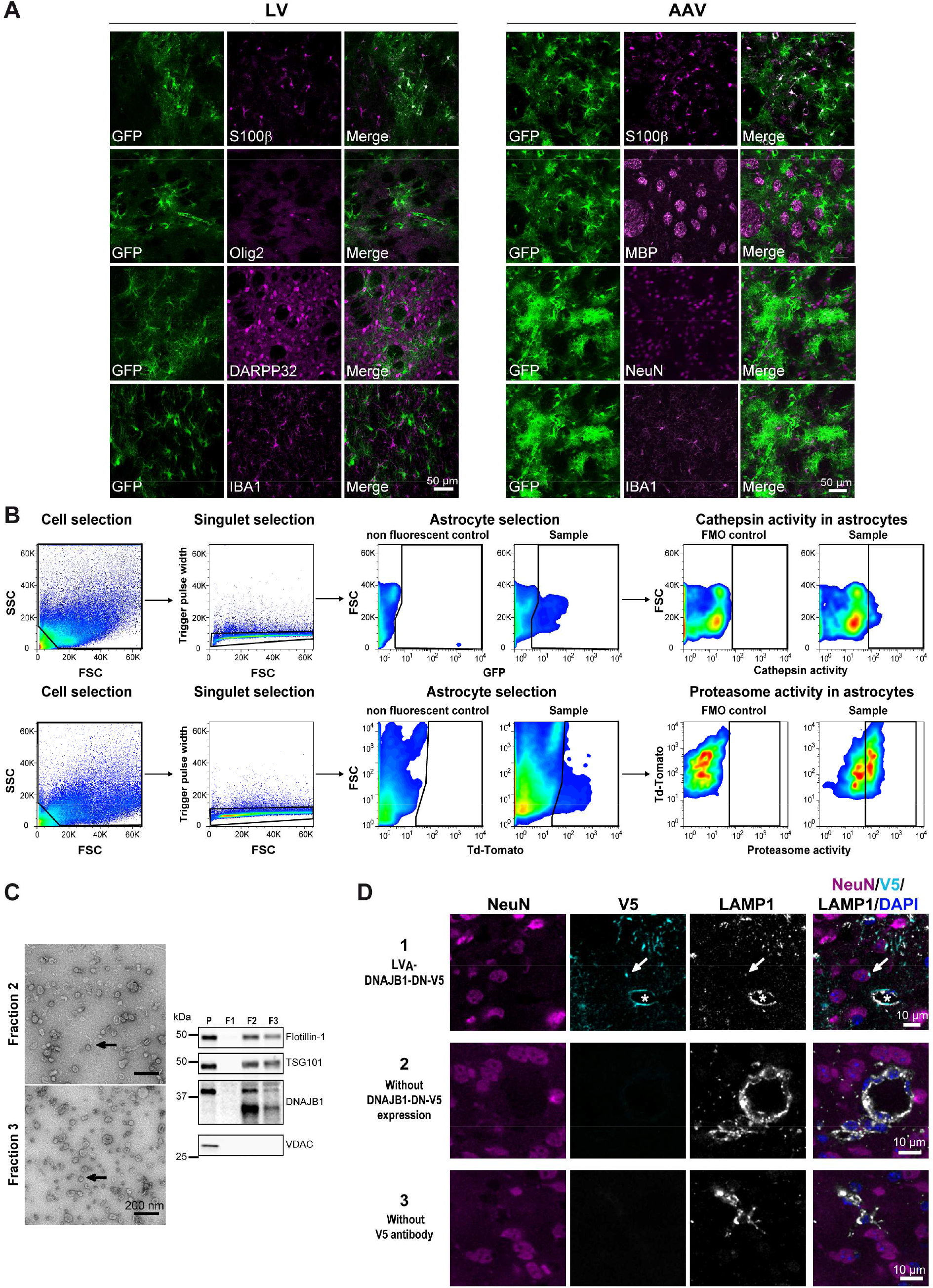
Validation of experimental procedures A. Validation of astrocyte transduction by viral vectors. After injection of LV or AAV encoding GFP and targeting astrocytes in the mouse striatum, GFP^+^ cells (green) co-express the astrocytic markers S100β, but not IBA1, NeuN, DARPP32, MBP or Olig2 (magenta), which are specific markers for microglia, neurons, oligodendrocyte or cells of the oligodendrocyte lineage respectively, demonstrating selective astrocyte tropism of both vectors. **B**. **Gating strategy for cathepsin and proteasome activity measurement on dissociated cells (in relation to** Fig. 6A**-C)**. Cells were gated on a side scatter/forward scatter plot, singlets were selected and then GFP^+^ or Td-Tomato^+^ astrocytes were gated based on a control non-fluorescent sample. Finally, the percentage of GFP^+^/cathepsin^+^ or Td-Tomato^+^/Proteasome^+^ astrocytes was quantified in each mouse after setting the gates on a sample processed similarly without the fluorescent activity probe (“Fluorescence Minus One” control). **C. Purity of the exosomal fractions (in relation to** Fig. 7F). Transmission electron microscopy of fractions 2 and 3 (F2, F3) obtained from the striatum of control mice evidences many circular vesicles with the typical size and shape of exosomes (arrow). Immunoblotting confirms that Flotillin-1 and TSG101, two exosomal proteins are enriched in F2 and F3, while the Voltage-Dependent Anion Channel (VDAC), a mitochondrial protein, is only found in the first pellet (P) of total/brain homogenate. DNAJB1 is abundant in exosomal fractions. **D. Controls for V5 immunostaining (in relation to** Fig. 8C**). D1,** Hdh140 mice injected as depicted in Fig. 8A to express DNAJB1-DN tagged with V5 (cyan) in astrocytes, display rare V5^+^ vesicles (arrows) in NeuN^+^ neurons (magenta). These vesicles are counterstained with the endo-lysosomal marker LAMP1 (white). DAPI is shown in blue. Note fine V5^+^ astrocyte processes that infiltrate the neuropil and send endfeet around a blood vessel (star). They display high LAMP1 expression. **D2,** On the contrary, no V5 staining is detected in brain sections from Hdh140 mice that do not express this transgene. **D3**, Omission of the primary antibody against V5 also results in an absence of V5 signal.

**Supplementary figure 2.**
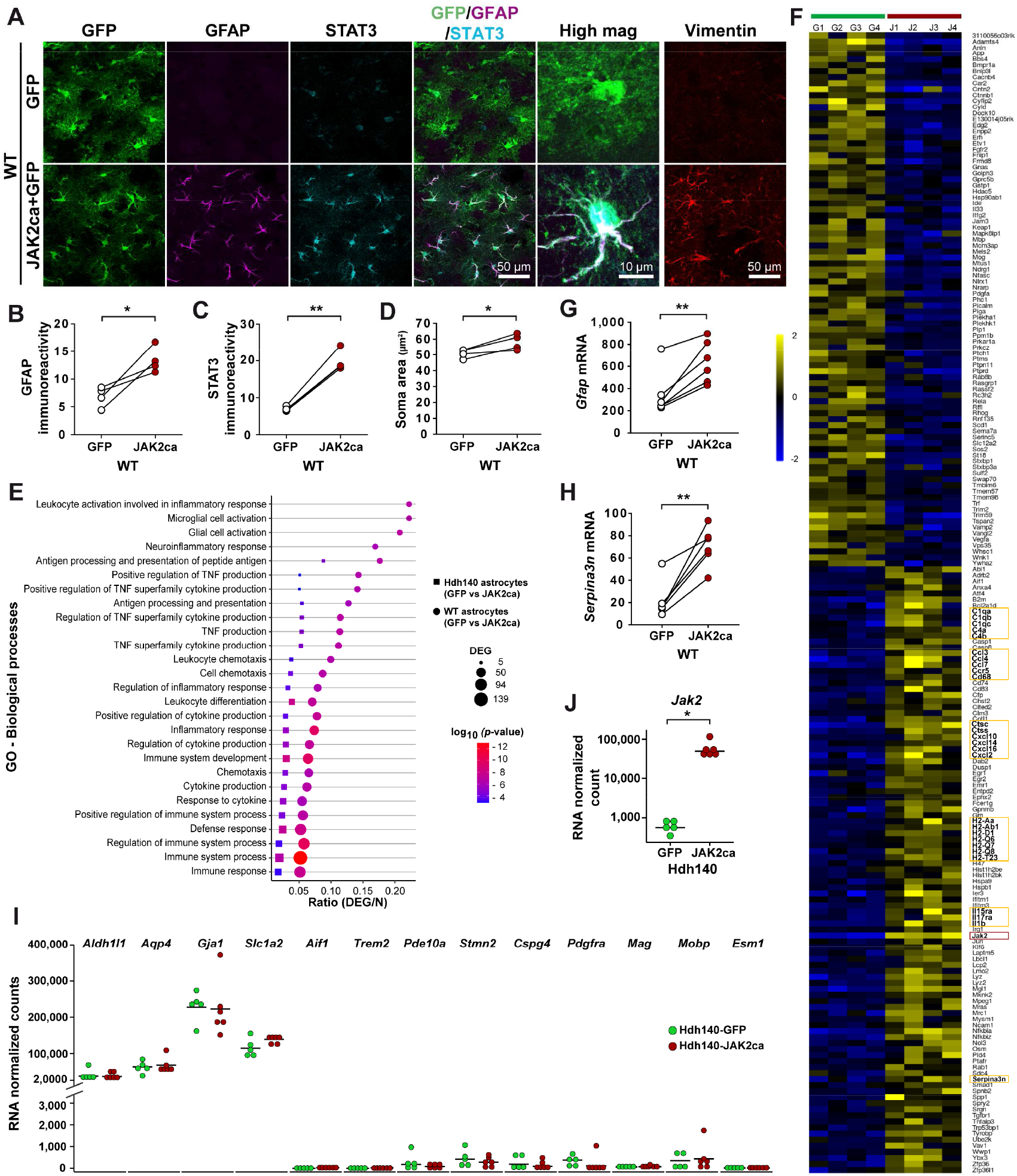
JAK2ca induces reactive changes in striatal astrocytes. A-H. Mice were injected as in Fig. 5A. **A-D.** Representative confocal images showing brain sections stained for GFP (green), GFAP (magenta), STAT3 (cyan) and vimentin (red) in WT- GFP and WT-JAK2ca mice (**A**). JAK2ca increases GFAP and vimentin immunoreactivity, triggers STAT3 accumulation in the nucleus and induces soma hypertrophy in astrocytes (quantification in **B, C, D**). **E,** GO analysis of differentially expressed genes (DEG) in WT- JAK2ca and Hdh140-JAK2ca sorted astrocytes identifies several biological processes linked to immunity and inflammation in both studies. **F.** In particular, gene expression heatmap for the four WT-GFP (G) and the four WT-JAK2ca (J) astrocyte samples shows that JAK2ca induces the expression of many genes involved in immunity and inflammation such as cytokines and chemokines, antigen presentation molecules and complement factors (highlighted in yellow boxes). Note that *Jak2* is also upregulated (red box). Color scale represents mean-centered expression (log2-transformed). **G, H**. JAK2ca-induction of *Gfap* (**G**) and *Serpina3n* (**H**) mRNA is validated by qPCR on bulk striatal samples prepared from WT-GFP and WT-JAK2ca mice. **I, J.** Mice were injected as in Fig. 5E. Sorted astrocytes from both Hdh140-GFP and Hdh140- JAK2ca mice express high levels of several astrocyte specific genes, while several markers of other cell types are barely detectable. **J.** *Jak2* mRNA are increased 100 fold in JAK2ca- astrocytes. **B-D,** Paired *t*-test; *n* = 4/group. **G, H**. Paired *t*-test; *n* = 6/group. **I-J**. *n* = 5-6/group.

## Supplemental tables

**Supplementary table 1:** **STAT3 involvement in gene regulation in Huntington’s disease astrocytes.**

| Tool | STAT3 rank among all significant transcription factors | p value<br>(provided by each tool) |
| --- | --- | --- |
| ChEA3-Enrichr | 53/1122 | 1.69E-22 |
| ChEA3-ENCODE | 96/104 | 2.11E-2 |
| ChEA3-ReMap Atlas | 11/123 | 1.91E-5 |
| HOMER | 25/55 | 1.00E-9 |
| TRANSFAC | 46/75 | 2.53E-6 |

STAT3 is identified among potential upstream transcription factors that regulate gene expression in Huntington's disease astrocytes. Data from Al-Dalahmah *et al.* 5.

**Supplemental table 2:** **Information on human samples**

| Group | Cause of death | Sex | Age<br>(y) | Post mortem delay<br>(h:min) | Brain weight (g) |
| --- | --- | --- | --- | --- | --- |
| Non-demented<br>control | Mediastinum carcinoma | Male | 45 | 08:50 | 1440 |
|  | Euthanasia (Pain) | Female | 55 | 07:30 | 1260 |
|  | Euthanasia (Pain) | Female | 60 | 05:30 | 1215 |
|  | Myocardial infarction | Male | 73 | 09:10 | 1500 |
| Huntington's<br>disease | Euthanasia | Female | 38 | 05:40 | 1140 |
|  | Euthanasia | Female | 57 | 06:40 | 1280 |
|  | Respiratory insufficiency | Female | 64 | 05:00 | 975 |
|  | Euthanasia | Male | 71 | 04:25 | 1320 |

**Supplemental table 3:**
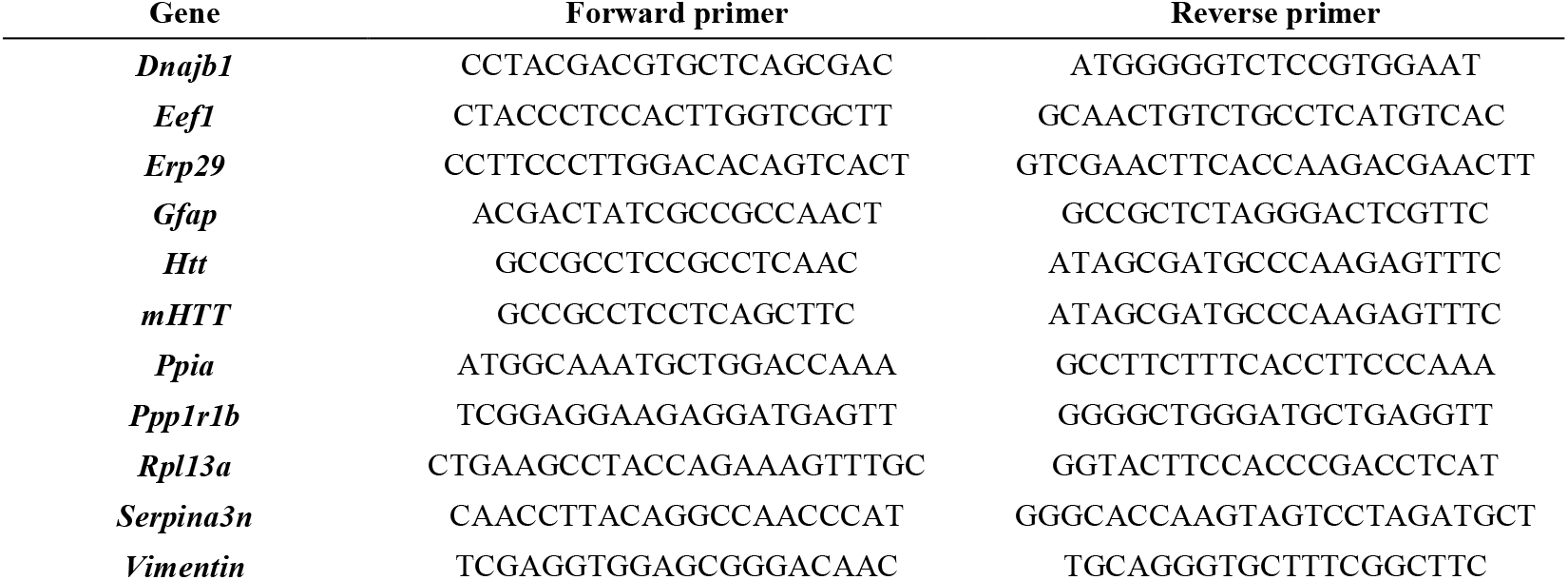
**Sequences of RT-qPCR primers**

## Supplementary Materials and Methods

### Stereotactic injections

WT mice were anesthetised with an *i.p.* injection of ketamine (100 mg/kg) and xylazine (10 mg/kg). For Hdh140 mice, xylazine was replaced by medetomidine (0.25 mg/kg) and anaesthesia was reversed by a *s.c.* injection of atipamezole (0.25 mg/kg) at the end of the surgical procedure. Lidocaine (7 mg/kg) was injected subcutaneously at the incision site, 10 min prior to surgery. Mice were given paracetamol (1.6 mg/ml) in drinking water for 48 h after surgery. Viral vectors were injected in the striatum (coordinates from Bregma: anteroposterior: + 1 mm, lateral: +/- 2 mm; ventral: - 2.5 mm from the dura, with the tooth bar set at 0.0 mm). LV were diluted in PBS with 1% bovine serum albumin (BSA), at a total final concentration of 100 ng p24/µl. AAV were diluted in 0.1 M PBS with 0.001% pluronic acid, at a final total concentration of 2.5 * 10^9^ viral genome (VG)/µl. Diluted viral suspensions (2-3 µl depending on the cohort) were injected at a rate of 0.2 - 0.25 µl/min with a pump.

### Immunohistology

Mice were either killed with an overdose of sodium pentobarbital (180 mg/kg) and perfused with 4% paraformaldehyde (PFA) or by cervical dislocation and one brain hemisphere was rapidly dissected and drop-fixed in 4% PFA. Brains were post-fixed for 24 h in 4% PFA, cryoprotected in 30% sucrose solution and cut on a freezing microtome into 30 µm-thick coronal sections. Series of brain sections were stored at -20°C in an anti-freeze solution until used for immunostainings.

#### Immunofluorescence

Brain sections were rinsed in PBS for 3 x 10 min and were blocked in 4.5% normal goat serum (NGS), 0.2% Triton X-100 in PBS (PBST) for 1 h at room temperature (RT). Brain sections were incubated overnight at 4°C with the primary antibodies diluted in 3% NGS/PBST: anti-DARPP32-647 (1:1,000, Mouse; Santa Cruz Biotechnology, Santa Cruz, CA, #sc-271111 AF647), anti-GFAP-Cy3 (1:1,000, Mouse; Sigma, Saint-Louis, MO, #C9205), anti-GFP biotinylated antibody (1:500, Goat; Vector Laboratories, Burlingame, CA, #BA- 0702), anti-IBA1 (1:500, Rabbit; Wako, Richmond, VA, #019-19741), anti-LAMP1 (1:500, Rat; Thermofisher Scientific, Waltham, MA, #14-1071-82), anti-MBP (1:500, Rabbit; Sigma, #M3821), anti-NeuN (1:500, Mouse; Chemicon, Billerica, MA, #MAB377), anti-Olig2 (1:500, Rabbit, Millipore, Burlington, MA, #Ab9610), anti-S100β (1:500, Mouse; Sigma #S2532), anti-V5 (1:1,000, Mouse; Invitrogen, #R96025), anti-Vimentin (1:1,000, Chicken; Abcam, Cambridge, UK, #ab24525) and NeuroTrace 640/660 (1:250; Thermofisher Scientific, #N21483). For DNAJB1 staining (1:100, Rabbit; Enzo Life Sciences, Farmingdale, NY, Hsp40/Hdj1 antibody, #ADI-SPA-400), brain sections were pre-treated in 0.1 M Tris-HCl, pH 9 at 95°C for 30 min and NGS was replaced by 5% BSA in blocking solution and antibody diluent. For EM48 (1:200, Mouse; Merck, Kenilworth, NJ, #MAB5374), sections were blocked in 3% BSA, 2% NGS/PBST and incubated with primary antibody in the same solution for 36 h. STAT3 immunostaining was performed as described previously^1^. For V5/LAMP1/NeuN immunostaining, brain sections were pre-treated in 0.01M citrate buffer at 95°C for 30 min. Brain sections were then rinsed 3 x 10 min in PBS and incubated with appropriate secondary Alexa Fluor-conjugated antibodies (1:1,000, Goat; Invitrogen) or for GFP staining with Streptavidine-FITC (1:1,000, Thermofisher Scientific, #SA100-02) in 3% NGS/PBST for 1 h at RT. Brain sections were rinsed three times with PBS before being mounted on SuperFrost® Plus slides (ThermoFisher Scientific) and coverslipped with Fluorsave™ (Calbiochem, Darmstadt, Germany). For V5/LAMP1/NeuN immunostaining, dilutions of secondary antibodies were filtered with a 0.22µm filter and sections were incubated 5 min in the Autofluorescence Eliminator Reagent (Millipore, #2160), according to the instructions ollowing provided protocol,to reduce tissue autofluorescence, before rinsing and mounting with Vectashield antifade mounting medium. Double or triple immunofluorescent stainings were performed successively, with each antibody incubated alone.

#### Immunohistochemistry with EM48 or Ubiquitin antibodies

Brain sections were pre-treated in 0.3% H2O2, blocked in 10% NGS/PBST (Ubiquitin) or 3% BSA/PBST (EM48) and incubated overnight at 4°C in the same diluent, with primary antibodies directed against EM48 (1:200) or Ubiquitin (1:1,000, Rabbit; Dako, Santa Clara, CA, z0458). After rinsing, brain sections were incubated with biotinylated secondary antibodies (1:1,000, Vector Laboratories) for 1 h at RT. Finally, they were incubated with the Vectastain Elite ABC Kit (Vector Laboratories) and revealed with the VIP kit (Vector Laboratories). Sections were rinsed three times with PBS before being mounted on SuperFrost® Plus slides, dehydrated and coverslipped with Eukitt (Sigma).

### Staining of human brain sections

Frozen blocks of the putamen were obtained from the Netherland Brain Bank, Netherlands Institute for Neuroscience, Amsterdam (open access: www.brainbank.nl). All Material has been collected from donors from whom a written informed consent for a brain autopsy and the use of the material and clinical information for research purposes had been obtained by the Netherland Brain Bank. Four Huntington’s disease patients (Vonsattel stage III) and four control subjects matched for age, sex and post-mortem delay were analysed (see **Supplementary table 2**). Blocks were cut into 10 µm sections on a cryostat, post-fixed in 4% paraformaldehyde; for 1 h and in ice-cold methanol for 10 min. Sections were then incubated in 0.1 M Tris-HCl (pH 9) at 95°C, for 20 min, in 0.3% H2O2 for 30 min, and in 5% BSA/PBST for 1 h. Sections were incubated 48 h at 4°C in 5% BSA/PBST, with primary antibodies against GFAP (1:10,000, Rabbit, Dako, #Z033429-2), STAT3 (1:200), NeuN (1:2,000) or DNAJB1 (1:100). After rinsing, sections were incubated with biotinylated secondary antibodies (1:1,000, Vector Laboratories) for 1 h at RT, and after rinsing, with the Vectastain Elite ABC Kit for 1 h and revealed with the DAB substrate kit (Vector Laboratories). Sections were incubated for 2 s in 50% Mayer’s hemalum solution (Merk, #109249) and rinsed with tap water before being dehydrated and coverlslipped with Eukitt. Representative images in each panel were taken with a Leica microscope, in the same anatomical region identified on consecutive sections.

### Protein extraction

Mice were killed by cervical dislocation and their brains were rapidly collected. The striatum was dissected out on ice, snap frozen in liquid nitrogen and stored at -80°C until protein extraction. Samples were homogenized by sonication in twenty volumes of lysis buffer [50 mM Tris-HCl pH 8, 150 mM NaCl, 1% Triton X-100 (Tx) with 1:100 phosphatase inhibitors (Sigma, cocktail 2) and 1x protease inhibitors with EDTA (Roche, Basel, Switzerland)] and centrifuged at 16,000 g for 30 min at 4°C. The supernatant contains Tx-soluble proteins and was used for immunoblotting. The Tx-insoluble pellet was washed with PBS and centrifuged at 16,000 g for 5 min at RT. The pellet was sonicated in a second lysis buffer [50 mM Tris-HCl pH 8, 2% Sodium Dodecyl Sulfate (SDS) with 1:100 phosphatase and protease inhibitors and centrifuged at 16,000 g for 30 min at 4°C. The supernatant was collected and used for immunoblotting, as the SDS-soluble fraction.

### Exosome isolation

Mice were killed by cervical dislocation and their brains were rapidly collected to isolate exosomes, as described in Vella et al.^2^. The striatum was dissected out on ice, snap frozen in liquid nitrogen and stored at -80°C until processing. Two or three striata were pooled by sample. Frozen striata were incubated in 75 U/ml collagenase III in Hibernate E solution (Thermofisher Scientific, 8 µl/mg) at 37°C under agitation. After 5 min, pieces of striatal tissue were gently pipetted up and down with a 1 ml pipette, then with a large diameter fire-polished Pasteur pipette and incubated at 37°C for 10 min. Then, tubes were gently inverted and returned to the incubation bath for 5 min (total incubation at 37°C: 20 min). Tubes were collected on ice and protease inhibitors were added at 1x final concentration. To discard debris, samples were sequentially centrifuged at 4°C at 300 g for 5 min, 2,000 g for 10 min and 10,000 g for 30 min. The last supernatants were collected and Hibernate E with protease inhibitors was added to a final volume of 3 ml. The first pellet was resuspended in 150 µl of 50 mM Tris-HCl, 1% SDS, 150 mM NaCl, 1 mM EDTA, pH 7.4 with protease inhibitors, and sonicated to be analysed by immunoblotting as total brain homogenates. A sucrose step was prepared for each sample as 0.3 ml of 2.5 M sucrose, 0.4 ml of 1.3 M sucrose, 0.4 ml of 0.6 M sucrose. The sample (3 ml in Hibernate E) was overlaid on top of the gradient. Sucrose steps were then centrifuged at 180,000g for 190 min at 4°C in a SW60 swinging rotor (Beckman, Brea, CA). After removing the top 2.6 ml of the step, three 0.4 ml-fractions (F1 to F3) were collected. Each fraction was diluted in 0.9 ml cold Dulbecco’s PBS (DPBS) with protease inhibitors and centrifuged at 100,000g for 1 h at 4°C on a fixed TL110 rotor (Beckman). The pellet containing vesicles was resuspended in 10 µl of cold DPBS with protease inhibitors and further diluted in loading buffer for immunoblotting or frozen until imaging by transmission electron microscopy (TEM).

The purity of exosomal fractions two and three was controlled by immunoblotting for known exosomal proteins (Flotillin-1 and TSG101) and for the mitochondrial protein voltage-dependent anion channel (VDAC), which is not present in exosomes (**Supplementary Fig. 1C**). These fractions were also analysed by TEM as described previously^3^. Exosomal fractions were mixed with an equal volume of 4% PFA and incubated for 20 min at 4°C. Fixed vesicles were then applied to carbon coated TEM grids and allowed to adsorb for 20 min at RT. TEM grids were then washed by sequential transfers on drops of PBS, incubated for 5 min at RT in 1% glutaraldehyde, and then washed by sequential transfers on drops of distilled water. Following negative-staining with 1% Gadolinium triacetate (EM stain 336, Agar Scientific, Stansted, UK) for 10 min at RT, samples were imaged in a Jeol 1400 transmission electron microscope (Jeol, Croissy/Seine, France). Images were recorded with a Gatan Orius CCD camera (Gatan, Pleasanton, CA) and processed with the Image J software.

### Immunoblot

Protein concentrations were measured by the bicinchoninic acid assay (Pierce, Waltham, MA). Samples were diluted in loading buffer with dithiothreitol (NuPAGE^®^ LDS sample buffer and sample reducing agent, Invitrogen). Proteins (10 µg for Tx-soluble fraction and 5µg for SDS-soluble fraction) were loaded on a 7.5% or 4-12% Criterion™ TGX Stain-Free Protein Gels (Bio-Rad, Hercules, CA). Protein concentrations in exosomal fractions were below detection level. Instead, an equal volume of each fraction was loaded on the gel. Migration was performed at 200 V for 30 min in Tris-glycine buffer (Bio-Rad) and proteins were transferred on a nitrocellulose membrane with the Trans-Blot Turbo™ Transfer System (Bio-Rad). After 3 x 10 min rinses in Tris Buffer Saline and 0.1% Tween 20 (TBST), membranes were blocked in 5% milk in TBST for 1 h at RT and incubated for 3 h at RT, or overnight at 4°C with the following primary antibodies: anti-Actin (1:5,000, Mouse; Sigma, #A2066), anti-Flotillin-1 (1:1,000, Rabbit; Sigma, #F1180), anti-DNAJB1 (1:1,000), anti-HTT (2B4, 1:1,000, Mouse; Millipore, #MAB5492), anti-mHTT (1C2, 1:1,000, Mouse; Millipore, #MAB1574), anti-Ubiquitin (1:1,000), anti-TSG101 (1:500, Rabbit; Sigma, #HPA006161), anti-α-tubulin (1:5,000, Mouse; Sigma, #T5168), and anti-VDAC (1:1,000, Rabbit; Abcam, #ab15895). After 3 x 10 min washes in TBST, membranes were incubated for 1 h at RT with HRP-conjugated secondary antibodies (1:5,000, Vector laboratories) diluted in TBST with 5% milk. Membranes were incubated with the Clarity Western ECL substrate (Bio-Rad) and the signal was detected with a Fusion FX7 camera (Thermofisher Scientific). Band intensity was quantified with Image J and normalized to actin or α-tubulin. For quantification of exosome proteins, the Stain-free technology (Bio-Rad) was used. Stain-free gels were exposed to UV light to activate tryptophan residues, resulting in UV-induced fluorescence of total loaded proteins. UV exposition and chemiluminescence acquisition were done with a ChemiDoc XRS+ system (Bio-Rad). DNAJB1 bands were normalized to total stained proteins using Image Lab Version 5.2.1 software (Bio-Rad). Each protein of interest was assessed at least on two different immunoblots.

### RNA extraction on bulk samples and RT-qPCR

Mice were euthanized with an overdose of sodium pentobarbital, their brains were rapidly collected, the GFP^+^ area was dissected out under a fluorescent macroscope (Leica) and lysed in Trizol with a MagNa Lyser instrument (Roche). Samples were stored at -80°C until further processing. Samples were placed 5 min at RT before addition of chloroform for 3 min, and centrifugation at 12,000 g for 15 min at RT. Aqueous phase was collected and 1 volume of 70% ethanol was added. Samples were transferred onto an RNeasyMin Elute spin column and RNA was purified according to manufacturer’s instructions, with on-column DNase treatment (RNeasy micro kit, Qiagen, Hilden, Germany). RNA was eluted with 14 µl of RNase-free deionized water and stored at -80°C before transcriptomic analysis. RNA quality and integrity was evaluated with an Agilent RNA 6000 Pico assay and the Agilent 2100 Bioanalyzer (Agilent technologies, Santa Clara, CA). Reverse transcription was performed with the VILO™ kit according to the manufacturer’s protocol (SuperScript^®^ VILO™ cDNA synthesis kit; Life Technologies, Carlsbag, CA). Samples were diluted in H2O with 100 µ g/ml BSA at 0.2 ng/µl and mixed with 250 nM of each primers and iTaq Universal SYBR Green Supermix (Bio-rad) for qPCR. PCR efficiency was between 85 and 110% for each set of primers (sequences shown in **Supplementary table 3**). Nuclease-free water and samples without reverse transcription were used as negative controls. Expression levels of transcripts of interest were normalized with the ΔCt method to the abundance of the best combination of normalizers among *Eef1*, *Erp29, Ppia, Rpl13a,* as identified with the Genorm method, implemented in Bio-rad CFX Manager software.

### Microarray analysis of acutely sorted astrocytes

WT mice injected with AAV-GFP or AAV-JAK2+AAV-GFP (same total viral load) were killed and their striatum rapidly collected in HBSS. The two striata of four mice were pooled before processing. Astrocytes were sorted as mentioned in section “*Quantification of cathepsin and proteasome activities in astrocytes*” in the main text; except that dissociated cells were prepared without the steps of myelin removal and probe incubation.

A total of ten sorted cell samples were processed for microarray analysis (four GFP^+^ astrocytes in the control WT-GFP group, four GFP^+^ astrocytes in the WT-JAK2ca group and two GFP^-^ cells in the WT-GFP group to validate astrocyte sorting efficiency). RNA was extracted as described in the previous paragraph. RNA quality and integrity was evaluated with an Agilent RNA 6000 Pico assay and the Agilent 2100 Bioanalyzer (Agilent technologies). RNA was amplified with the Ovation PicoSL VTA system V2 kit (NuGen technologies, San Carlos, CA). Single strand DNA and single primer isothermal amplification cDNA were purified with Agencourt RNAClean XP (NuGen technologies). cDNA concentration was measured with a Nanodrop-1000 spectrophotometer (Labtech France). The Encore biotiNL kit (NuGen technologies) was used for the fragmentation and labelling of the purified SPIA cDNA prior to hybridization on the Illumina BeadChip mouse WG-6v2, which contains more than 45,000 unique 50-mer oligonucleotides (Illumina, San Diego, CA). BeadChips were scanned on the Illumina Iscan. A control summary report was generated by the GenomeStudio software (Illumina) to evaluate the performance of built-in controls (variation in hybridization and background signals and background/noise ratio). Quantile normalization without background subtraction was applied to all samples within an analysis, with GenomeStudio software. First, to validate sorting efficiency, we compared the four GFP^+^ astrocyte samples with the two GFP^-^ cell samples from the WT-GFP group (*t* test, *p* < 0.05 and Fold change > 1.5). Then, we compared the 4 GFP^+^ astrocyte samples from WT-GFP mice with the 4 GFP^+^ astrocyte samples from the WT-JAK2ca mice to identify transcriptional changes induced by JAK2ca in astrocytes. To study only genes with reliable expression levels, we included probes with a “signal *p* value” above 0.01 in more than 50% of the samples. Samples with a signal *p* value below 0.01 were arbitrarily given a signal value of 75 (the lowest possible signal value being 76.5). Expression levels of detectable probes were compared between WT-GFP and WT-JAK2ca astrocytes with the R (v3.6) Bioconductor (v3.13) limma (linear models for microarray data, v3.44.3) package^4^. Principal component analysis was performed with the *plotPCA* function from R DESeq2 package (v1.12.3)^5^, also in Bioconductor. Log-transformed data were plotted using the top 500 most variable genes.

Analysis for enriched Gene Ontology (GO) terms [Biological Processes (BP), Molecular Functions (MF) and Cellular Component (CC)] and Kyoto Encyclopaedia of Genes and Genomes (KEGG) pathways were performed with an adapted function from the same R package limma in Bioconductor repository. Significant GO-BP and GO-MF entries relevant to proteostasis were selected, and all differentially expressed genes belonging to these GO entries were plotted as a network, with the *cnetplot* function from the R package DOSE (v2.10.6) from Bioconductor.

### RNAseq analysis of acutely sorted astrocytes from Hdh140 mice

To analyse the transcriptome of astrocytes acutely sorted by FACS from the striatum of 12- 14 month-old Hdh140 mice injected with AAV-GFP or AAV-JAK2ca (N=5-6), we followed the protocol described in Ceyzériat et al.^6^, except that the lower number of astrocytes sorted per sample was not compatible with RNA quality assessment, and full length double strand cDNA libraries were amplified with 16 LD-PCR cycles. The final RNAseq libraries were then sequenced on a HiSeq 2500 Illumina platform (2 × 100 bp). Quality control of sequencing data was performed with FastQC (v0.11.9)^7^. Reads were mapped on the GRCm38 (mm10) mouse genome assembly with Hisat2 (v2.2.1)^8^. Quantification of reads associated with genes was achieved with featureCounts (v2.0.0)^9^, and differential gene expression analysis was performed with DESeq2 (v1.28.1) Bioconductor (v3.13) package on R (v4.0.2)^5^. Only genes with a raw number of counts ≥ 10, in at least 3 samples were analysed. Results were considered statistically significant for an adjusted *p* value ≤ 0.1 and fold-changes ≥ 1.5. Principal component analysis, GO and KEGG analysis were performed on R as described for microarray data, and fast preranked gene set enrichment analysis (GSEA) was carried out with the fgsea (v1.16.0) package on R Bioconductor as well^10^.

Venn diagrams were generated with BioVenn (v 1.1.3) R CRAN package^11^ using all GO-BF, GO-CC and GO-MF identified as enriched in the DEG list of WT-JAK2ca and Hdh140-JAK2ca astrocytes and with more than 2 DEG per GO term.

### Identification of regulatory transcription factors

We performed bioinformatics analysis on the list of 2,250 differentially expressed genes between control and Huntington’s disease astrocyte nuclei isolated from the cingulate cortex of human subjects (Additional file 9 in Al-Dalahmah et al.^12^). To identify putative upstream transcription factors, we used HOMER^13^ and Pscan with the TRANSFAC database^14^, two tools based on motif recognition in the promoter region of regulated genes. We also used three of the assembled transcription factor libraries from publicly available data (e.g. chromatin immunoprecipitation experiments, co-expression datasets) offered by the ChEA3 tool^15^: Enrichr, ENCODE and ReMap.

### Magnetic resonance imaging and analysis

T2 anatomical imaging was performed on a horizontal 11.7 T Bruker scanner with a quadrature cryoprobe (Bruker, Ettlingen, Germany) for radiofrequency transmission and reception.

Six Hdh140 mice (17-18 month-old) injected in one striatum with LV-GFP and the other with LV-JAK2ca+LV-GFP (same total viral load) were anesthetised 2 months later by 3% isoflurane in 1:1 air/O2. Mice were positioned with mouth and ear bars in a dedicated unmagnetic bed. Mouse temperature was monitored with a rectal probe and maintained at 37 °C with regulated water flow. Respiratory rate was continuously monitored using PC SAM software (Small Animal Instruments, Inc., Stony Brook, NY, USA). Isoflurane level was adjusted around 1.5% to maintain a respiratory rate at 60-80 per minute.

High-resolution anatomical T2-weighted images were acquired in coronal orientation with Multi Slices Multi Echoes sequence as described in ^16^. The two striata were manually delineated on each 300 µm-slice on high-resolution images by asingle operator and volume was calculated for each animal according to the Cavalieri method.

GluCEST imaging was performed on the same magnet using a volume coil for radiofrequency transmission and a quadrature surface coil for reception (Bruker, Ettlingen, Germany). Three gluCEST images centered on the mid-striatum were acquired with a 2D fast spin-echo sequence preceded by a frequency-selective continuous wave saturation pulse, according to the procedures described in ^16^). GluCEST images were processed pixel-by-pixel and analyzed using in house MATLAB programs as described in ^16^. The striatum was manually delimited on the reference CEST image acquired without saturation to extract the percentage of gluCEST contrast for each striatum in each mouse.

